# Origins and timing of emerging lesions in advanced renal cell carcinoma

**DOI:** 10.1101/2021.06.27.450111

**Authors:** Andrew Wallace, Sima P. Porten, Amy A. Lo, Daniel Oreper, Nicolas Lounsbury, Charles Havnar, Ximo Pechuan-Jorge, Oliver A. Zill, Maxwell V. Meng

## Abstract

**Purpose:** Renal cell carcinoma (RCC) with venous tumor thrombus (VTT) arising from the primary tumor occurs in 4-10% of cases and is associated with advanced disease. RCC with VTT and distant metastasis represents a unique clinical entity, and provides opportunities to examine the origins and relative timing of tumor lesion emergence and to identify molecular correlates with disease state.

**Experimental Design:** We performed genomic and evolutionary analyses on 16 RCC patients with VTT, with eight also having metastases, using multi-region exome and RNA sequencing.

**Results:** No genomic alterations were specifically associated with the VTT or metastasis lesions; each tumor had multiple hallmark driver alterations, consistent with advanced disease state. We found that 21% (3/14) of clear-cell RCC cases could be assigned a previously defined “evolutionary subtype”. Somatic mutation signatures were largely consistent with previously established RCC signatures, and showed low heterogeneity across regions of each tumor. Mismatch repair and homologous recombination (“BRCA-ness”) deficiency signatures consistently co-occurred across most tumors, suggesting a pervasive role for intracellular DNA damage in RCC and the potential for related treatment strategies. Phylogenetic timing analysis of metastatic cases suggested that in most tumors, metastases branched from the primary tumor prior to formation of VTT and in some cases before diversification of the primary tumor. Both VTT and the earliest metastases were predicted to emerge many years prior to diagnosis. Transcriptional landscape analysis identified key differences distinguishing each lesion type from primary tumor: VTT upregulated TNF*α* signaling and associated inflammatory pathways, whereas metastases upregulated *MTOR* signaling.

**Conclusions:** Our results provide a map of how RCC tumors can evolve, with metastatic clones typically emerging early in RCC development and taking hold via *MTOR* signaling, and later formation of VTT via local inflammatory processes.

**Statement of Translational Relevance:** Renal cell carcinoma (RCC) is a deadly and relatively common malignancy, which often presents as or progresses to metastatic disease. We used multi-region sequencing of RCC patients with venous tumor thrombus (VTT) and metastasis to ask how and when new lesions arise from the primary tumor, and what genomic factors contribute to their spread. Phylogenetic analysis of patients with VTT and co-presenting metastases suggested that in most cases, the VTT and metastases derive from distinct tumor clones. Moreover, metastatic clones often appear many years prior to diagnosis. We found that local TNF*α* inflammation may contribute to VTT formation, whereas MTOR signaling is associated with metastases. Our study sheds light on the relationship of VTT and metastases, suggests therapeutic and biomarker strategies for RCC, and points to the need for early detection studies in RCC to better understand when metastases emerge and to identify at-risk patients.

## Introduction

Renal cell carcinoma (RCC) is the most common type of kidney cancer, and is one of the most common neoplasms worldwide (1). RCC exhibits local vascular invasion in up to 10% of cases, and the resulting venous tumor thrombus (VTT) impacts treatment options and may present challenges for surgical intervention (2,3). Recent work has made strides towards genomic characterization of clear cell RCC (ccRCC) tumors with VTT, and has begun to delineate the genomic features of VTT tumors from non-VTT tumors (4,5). For example, ccRCC tumors with VTT exhibit greater tumor mutational burden, higher genomic instability, and higher tumor proliferative indices than tumors without VTT (4,6).

Several studies have employed multi-region sequencing (MR-seq) to explore patterns of tumor evolution in RCC for cases both with and without VTT (5–9). RCC tumors exhibit branched evolution leading to high intratumoral heterogeneity (ITH) and substantial subclonal diversity. MR-seq has also shed some light on the evolutionary patterns that give rise to VTT. Warsow et al. (5) demonstrated that while multiple VTT regions typically derive from a single common ancestor, they likely originate from subclones common to primary tumor regions.

Turajlic et al. (6) found similar patterns, with the VTT seemingly derived directly from the most recent common ancestor of all tumor clones in the greatest proportion of assayed cases. However, the relationship between VTT and metastases remains unclear. Many patients with VTT present with metastasis (20-50%), but VTT formation and metastatic spread may not necessarily be linked (10,11). Broadly, it is possible that either metastases could be seeded from the VTT clone via hematogenous spread, or that metastases are derived from separate clones that may originate earlier or later than the VTT clone. As VTT physically emerge from the primary tumor, their genomic composition provides a point of comparison for the relative timing and subclonal origin of distant metastasis from the primary tumor.

The timing of metastatic spread is an issue of substantial clinical significance (12), and has been explored in other cancers such as CRC (13,14). Many patients with RCC present with metastases at the time of diagnosis (20-30%), however in some patients metastases only become evident over time (15). Knowledge of the relevant timelines is critical to properly inform screening regimens, as patients at risk for early metastasis should be subject to more regular screenings, and may impact treatment decisions. Therefore, quantitative approaches for estimating the timing of metastasis in RCC from genomic evidence are increasingly critical for improved prognosis, treatment algorithms, and ultimately clinical outcomes (16). Similarly, understanding which biological pathways contribute to the formation of new lesions and their spread from the primary tumor is important to inform therapeutic development and treatment decisions. Toward these ends, we performed MR-seq analysis of RCC with VTT and metastases--including both mutation and gene expression characterization--to estimate the timing of their clonal emergence and to identify biological pathways associated with each type of lesion. We further hypothesized that the resulting phylogenomic patterns might shed light on the clinical pictures of these patients and inform RCC treatment more broadly.

## Materials and Methods

### Clinical sample identification and pathology analysis

Institutional Review Board approval was obtained from the University of California San Francisco. Cases clinically diagnosed as renal cell carcinoma with venous tumor thrombus that contained banked tissue for research were identified. All patient identifiers were reassigned to protect anonymity. A surgical pathologist reviewed all slides associated with each case, established that primary tumor, metastatic tumor, and thrombus regions were available based on the gross description and confirmed diagnoses based on morphologic and immunohistochemical findings. Viable tumor content (% viable tumor/total epithelial surface area), stromal abundance (low, moderate or high) and presence of cystic change were estimated for each case by an independent pathologist. Retrospective chart review was performed to identify and capture relevant clinical and demographic information (Figure S1). A full cohort description and patient metadata are available in Tables S1 and S2.

### IHC analysis

IHC was performed on 4um thick FFPE tissue sections mounted on glass slides. IHC for PD-L1 clone SP263 (Roche Tissue Diagnostics, Tuscan, AZ, cat 790-4905) was performed on the Ventana Benchmark XT platform. The slides were pretreated with CC1 for 64 min followed by primary antibody incubated for 16 minutes at 37°C. The antibody was detected with the OptiView DAB IHC Detection Kit. PanCK and CD8 duplex chromogenic IHC was performed using established methods on the Ventana Discovery Ultra. The fraction of viable tumor cells (%) that express membrane PD-L1 were quantified. The overall immune phenotype was classified as “desert”, “inflamed”, “excluded” (two types: invasive margin or intratumoral) based on the predominant (>10%) location of CD8 positive cells in relation to the tumor. If no single pattern occurred in the majority of the tumor, then the slide was categorized as “heterogeneous”. Automated slide assessment was performed quantitatively using Visiopharm analysis software. Tissue area that was positive for the panCK was used to generate an epithelial tumor mask and the relative surface area of CD8 positive cells within stromal and epithelial tumor compartments was determined using Visiopharm software applications.

### Generation and processing of sequencing data

Exome sequencing libraries were created with Agilent SureSelectXT. SureSelect All Exon V6 bait probes were used to perform targeted capture at Q^2^ Solutions (Valencia, CA). Exome sequencing coverage was approximately 75 million 100bp paired-end reads, yielding an average depth (before removing duplicate reads) of 150X per sample. RNA-seq libraries were generated using the RNA Access platform. Sequencing coverage was approximately 50 million 50bp paired-end reads per sample.

Exome FASTQ files were trimmed to remove adapter contamination and low-quality trailing sequences, and then mapped to the GRCh38 reference genome with GSNAP (17). Mutations were called using both Lofreq2 (18) and Strelka (19), with the union of both outputs retained prior to further filtering. The filtering required mutations to have a minimum VAF of 0.05 and coverage of 20 in at least one sample in which they were called. Further, they were required to exhibit a maximum VAF of 0.01 in the normal sample, and a maximum ExAC (20) global frequency of 0.01. Somatic copy number alterations (SNCA) were identified using TitanCNA (21). RNA-seq FASTQ files were trimmed to remove adapter contamination and low-quality trailing sequences, and then mapped to GRCh38 using GSNAP (17).

### Driver alterations and mutational signature analysis

Driver alterations were identified as putative loss-of-function mutations in previously reported ccRCC and pRCC driver genes, including *VHL, PBRM1, SETD2, PIK3CA, MTOR, PTEN, KDM5C, CSMD3, BAP1, TP53, TSC1, TSC2, ARID1A, TCEB1, MET, NF2, KDM6A, SMARCB1, FAT1, STAG2, NFE2L2* and *CDK2NA,* as well as previously established arm-level driver SCNAs in chromosomes 3p loss, 5q gain, 4q loss, 8p loss, and 14q loss (7,22–24). Loss-of-function mutations were defined as indels, nonsense SNVs, splice site dinucleotide SNVs, and putatively deleterious missense SNVs according to SIFT (25), PolyPhen (26), or CONDEL (27).

The sample frequencies of SNV trinucleotide motifs were extracted using the SomaticSignatures R package (28). Signatures previously identified by Alexandrov et al (29) were then fit to the resulting motif spectra using non-negative least squares with the MutationalPatterns R package (30).

### Differential gene expression analysis

RNA-seq reads within gene coding regions were quantified using HTseq (31), and differential expression was assessed using DESeq2 (32). Genes were considered differentially expressed if the comparison had p_adjusted_ ≤ 0.05, and the sign of the log-fold change was consistent among individual regions in more than half of the patients. The relevant pairwise comparisons were performed while controlling for patient-specific expression. Gene set overlap analysis of differentially expressed genes was performed using Enrichr (33), which performs a hypergeometric test, and gene set enrichment analysis (GSEA) was performed using the R package fGSEA (34). For the latter, all genes included in the differential expression testing were ranked by Wald statistic of the comparison.

### Phylogenetic analysis and lesion emergence timing

Phylogenetic trees were constructed using the method of maximum parsimony, on the basis of binary presence or absence of each mutation per sample. Trees were reconstructed 100 additional times on nonparametric bootstrap samples of the input mutation sets, according to the method of Felsenstein (35). We used the phangorn (36) and the APE R packages (37) for tree construction and bootstrapping respectively. To assess the relative order of region emergence, the sums of the internal branch lengths from the root to the region parent nodes were compared in each bootstrap replicate. In order to estimate absolute timing of lesion parent clone emergence, we adapted the linear mixed modeling approach of Mitchell et al (38), using the lme4 package (39). (See Supplementary Methods for further details.)

## Results

### Genomic alteration patterns suggest advanced disease, minimal overlap with previously described evolutionary subtypes

We investigated a cohort consisting of 16 total patients, 14 of whom had ccRCC, and 2 had pRCC (Figure S1, Table S1). A total of 78 distinct tumor regions from these patients were assayed for this study, including multiple primary regions and at least one VTT region from all patients, as well as at least one metastasis from eight of the 11 metastatic cases (Table S2).

Five patients had more than one metastasis sampled. Exome sequencing was performed on all patients and tumor regions along with one matched normal sample per patient. RNA sequencing (RNA-seq) was performed on 56 tumor regions from 12 patients along with one matched normal sample per patient. Somatic point mutation analysis identified a range of 3.6-25.2 unique mutations/Mb per patient, and a range of 1.8±0.24 mutations/Mb to 8±2.4 mutations/Mb per sample (Table S3). Tumors in this cohort exhibited a wide range of global clonality--the percentage of mutations in any given sample that were detected in all samples from the same tumor--with values as low as 6.5% and as high as 81% (Table S3). The ccRCC cases exhibited a median average of 48% globally clonal mutations, higher than previous reports have indicated (8). Both pRCC patients had an average of 73% globally clonal mutations, higher than all but two non-metastatic ccRCC cases (73% in PT6 and 81% in PT13). Lower intratumoral heterogeneity and higher global clonality have been previously reported for pRCC, when compared with ccRCC (40).

We analyzed the driver alteration landscape of all patients and samples, accounting for both somatic genomic alterations and gene expression of known driver genes in RCC (for details see Methods). All patients exhibited clonal, putatively inactivating mutations in at least one previously identified RCC driver gene (Figure 1A). Expression of canonical RCC driver genes was generally consistent with the expected behavior for tumor suppressors (downregulated in tumor relative to matched normal), with the exception of *FAT1*, *MET*, *SMARCB1*, and *TP53* (Figure S3). Driver mutations with at least 5% prevalence in this cohort displayed a surprising level of homogeneity relative to other RCC cohorts. In six patients these mutations were entirely clonal (i.e., were observed in all regions; PT7, PT8, PT10, PT11, PT13, PT16), and in an additional four patients there was only one region displaying heterogeneity (mutation presence/absence; PT5, PT6, PT14, PT15). There were several examples of parallel mutations in separate tumor regions of six patents (Figure 1B), consistent with previous studies, and suggesting selective pressure for loss of function in these genes (8).

**Figure 1.**
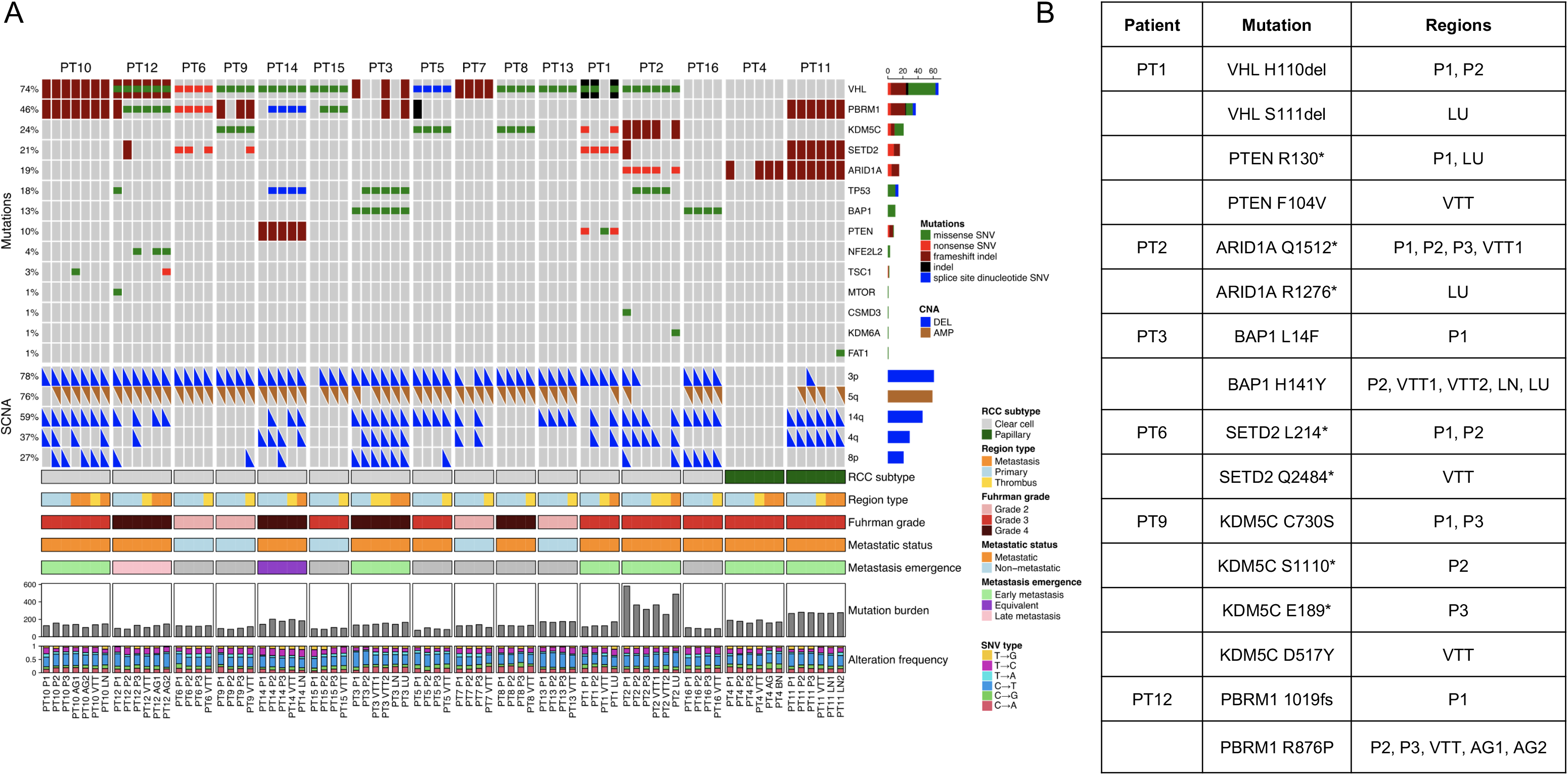
Genomic alteration landscape across tumor regions in 16 RCC patients with VTT. (A) An oncoprint showing the most prevalent somatic mutations and SCNAs found in this cohort. Prevalences of each alteration are indicated at left. Tumor lesion labels: P = primary tumor; VTT = venous tumor thrombus; AG = adrenal gland metastasis; LN = lymph node metastasis; LU = lung metastasis; BN = brain metastasis. (B) Parallel mutations found in distinct regions of six tumors.

All ccRCC patients exhibited either truncal *VHL* mutation or truncal loss of chromosome 3p, with co-occurrence of these truncal driver alterations in 10 patients. Consistent with previous observations (22), *PBRM1* was the second most commonly mutated of the assessed established driver genes, harboring globally clonal mutations in three ccRCC patients, and subclonal mutations in a further five. *PBRM1* mutation has been associated with better prognosis, except when *BAP1* is co-mutated, which is associated with aggressive disease and poor prognosis (41,42). However, PT3 had *PBRM1, BAP1* co-mutation but exhibited no evidence of disease on final follow-up (Figure S1). For the two papillary cases, PT11 exhibited mutations in a number of ccRCC driver genes (*PBRM1*, *SETD2*, and *ARID1A*), whereas patient PT4 exhibited only subclonal mutation of *ARID1A*. In metastatic patients, we observed an enrichment of SCNA losses in chr14q, 4q, and 8p relative to non-metastatic patients, consistent with the more advanced disease state. We did not identify any recurrently mutated genes associated with the VTT samples, aside from genes known to be frequent false positives due to long coding sequences or a large number of paralogs (Table S4).

To investigate whether these tumors followed prescribed patterns of evolution, we used canonical driver alterations to assign the ccRCC patients to the seven evolutionary subtypes described by Turajlic et al (7). We note that we were unable to consider the methylation status of driver genes; only loss-of-function mutations in driver genes and driver SCNAs were included. We were able to assign three of the 14 ccRCC cases confidently to evolutionary subtypes: PT6, PT9, and PT10 (Table 1). Four additional cases exhibited similarities to the established subtypes, but could not be clearly assigned. The remaining seven cases were inconsistent with the pattern of driver mutations characteristic of any of the seven subtypes.

**Table 1.**
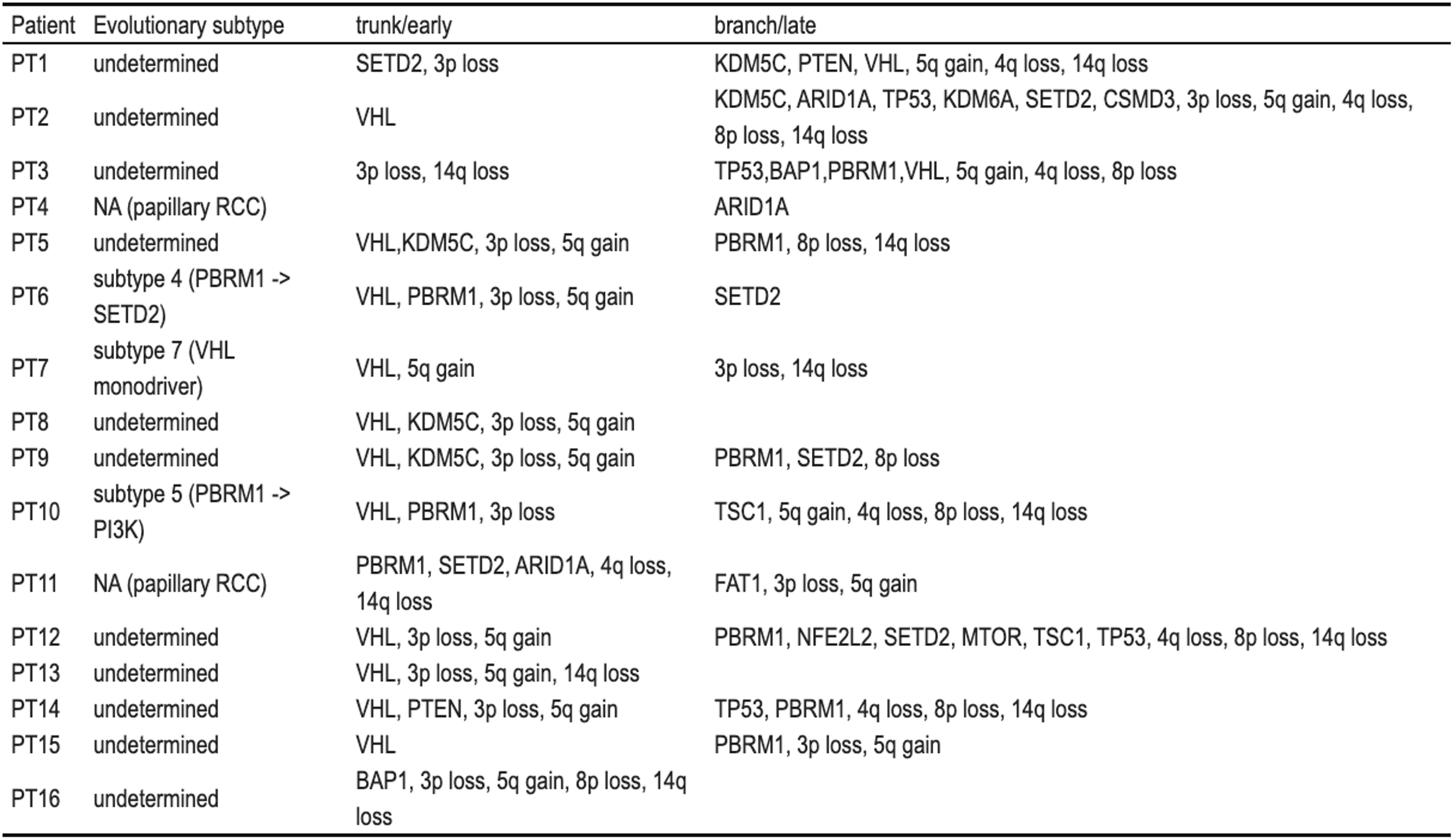
Phylogenetic placement of driver alterations (early/late) and concordance of each tumor with previously described evolutionary subtypes (6).

### Prevalence and co-occurrence of two types of DNA repair deficiency mutational signatures

We next asked whether specific mutational signatures were identified in patients with VTT. We examined somatic SNV substitution patterns, along with their trinucleotide context, and compared the resulting spectra to previously identified mutational signatures (29). Determination of the relative contribution of known signatures to the observed spectra revealed broad similarities within patients and heterogeneity among patients (Figure 2A). Extraction of the signatures predominantly contributing to each sample’s mutational spectra revealed an enrichment of signatures associated with age (S1A and S1B), mismatch repair deficiency (S6), and *BRCA1/2* mutation (S3) across a majority of patients (Figure 2B).

**Figure 2.**
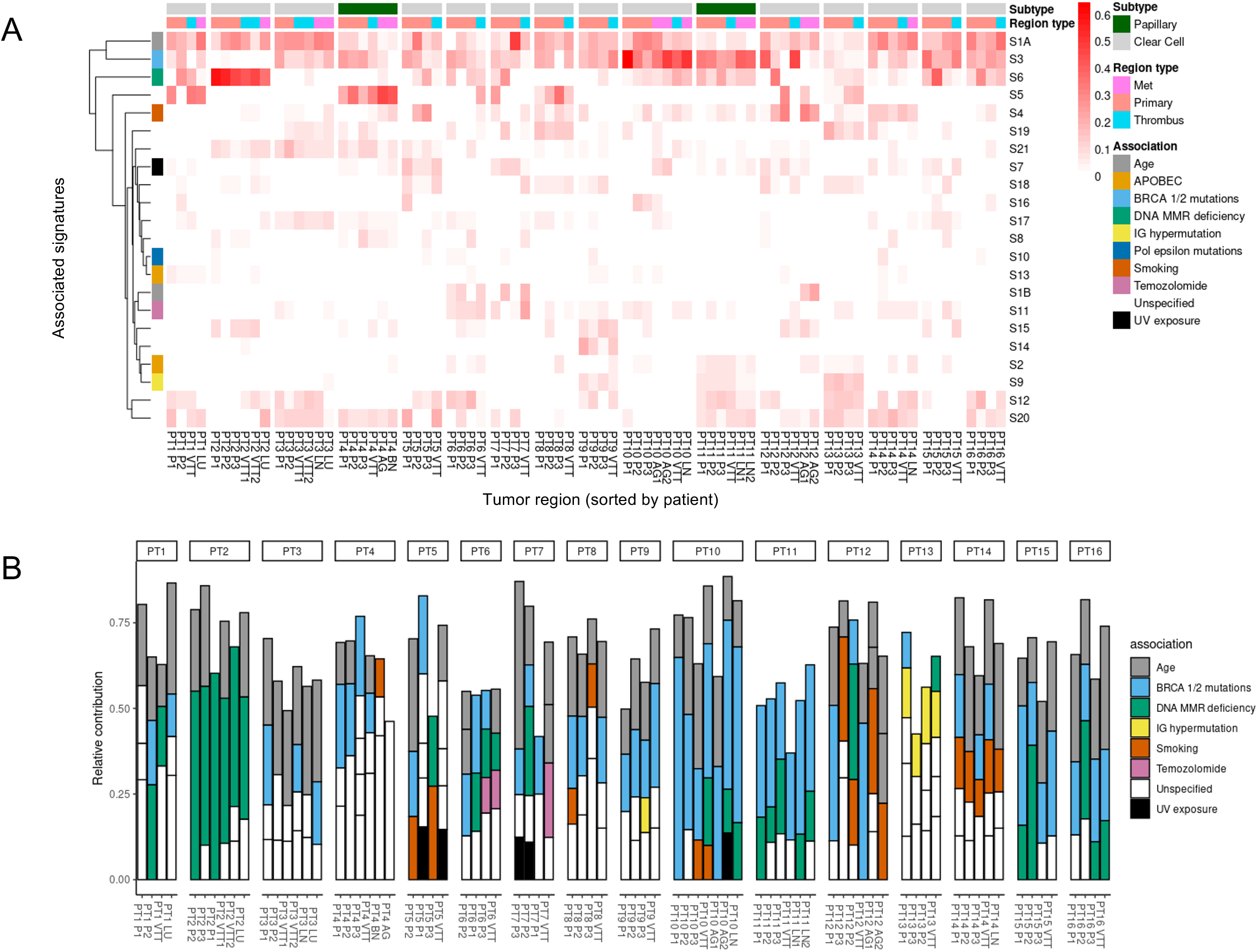
SNV mutational signature analysis. (A) Heatmap of previously established SNV mutational signatures to the mutational spectra of each tumor region across all patients. (B) Contribution of previously established SNV mutational signatures to the mutational spectra of each region across all patients. Included signatures are restricted to those with >10% contribution in a sample.

The most prevalent signatures across our patients were signatures S3 and S1A. S3 is typically associated with *BRCA1*/*2* mutation and has been previously observed in RCC (5). Almost all patients (15/16) exhibited some degree of signature S3 (53/78 regions exhibited >10% contribution), but there were only two patients with potential deleterious *BRCA1/2* alterations. Patient PT8 exhibited a truncal frameshift variant in *BRCA1*, while patient PT4 exhibited a truncal variant in a splice region of the gene *ZAR1L*, which occurs immediately upstream of *BRCA2*. The presence of signature S3 in the absence of BRCA mutations is a described phenomenon (43,44). The only case lacking the S3 “BRCA-ness” signature was patient PT2, who instead displayed a strong signature S6 (associated with DNA mismatch repair deficiency). Strikingly, this patient also harbored a truncal nonsense mutation in *MLH1*, which is known to cause mismatch repair defects but is rare in RCC (45). Moreover, this patient exhibited by far the most mutations per sample (7.9 mutations/Mb compared to an average of 2.8 mutations/Mb for the other patients), and by far the most total unique mutations (25.2 mutations/Mb compared to an average of 5.7 for the other patients). Signature S6 was also identified as exhibiting >10% contribution to at least one tumor region in nine additional patients. This observation is consistent with previous reports, but is unexpected in light of the relatively low mutational burden of RCC (29). Interestingly, S3 and S6 co-occurred in at least one region in 14/16 patients, and co-occurred at >10% contribution each in eight patients. Signature S1A is associated with age, and also occurred in most patients (14/16) and in most tumor regions (>10% contribution in 55/78 regions). Presence of age-associated signatures is consistent with previous assessments of RCC mutational processes (29).

### Phylogenetic and timing analyses suggest relatively early emergence of metastasizing clones in some patients

We next asked whether the VTT acts as a reservoir for hematogenous metastases. If this were the case, VTT subclones should typically emerge prior to metastases and the metastases should share a clonal ancestry with the VTT. To test this hypothesis, we performed phylogenetic analysis using binarized somatic mutation calls for the eight patients that had primary, VTT, and metastasis lesions (Figure 3). In seven of eight patients, the VTT had a distinct clonal ancestry from the earliest metastasis (PT10, PT2, PT3, PT4, PT11, PT1, PT12). In PT14, the VTT and LN metastasis shared an immediate ancestor. PT10 was a more complex case, wherein the VTT shared an immediate ancestor with the LN metastasis, but the two adrenal gland metastases arose earlier from a distinct ancestor. To establish the certainty of the phylogenetic reconstruction, and in particular to establish the confidence in the placement of specific nodes, we performed non-parametric bootstrapping of the mutational tree (13). Overall across the eight patients, the prior assertions of distinct clonal origins for VTT and metastasis were well supported, with nodal bootstrap values consistently above 70 (top panels of Figure 3A-C). However, PT4 had lower bootstrap values for the two key nodes (44 and 41), suggesting less certainty in the phylogenetic placement of the clonal ancestors for the AG, BN, P1, P2, and VTT regions. These results suggest that in most patients the VTT likely did not act as a reservoir for hematogenous metastases.

**Figure 3.**
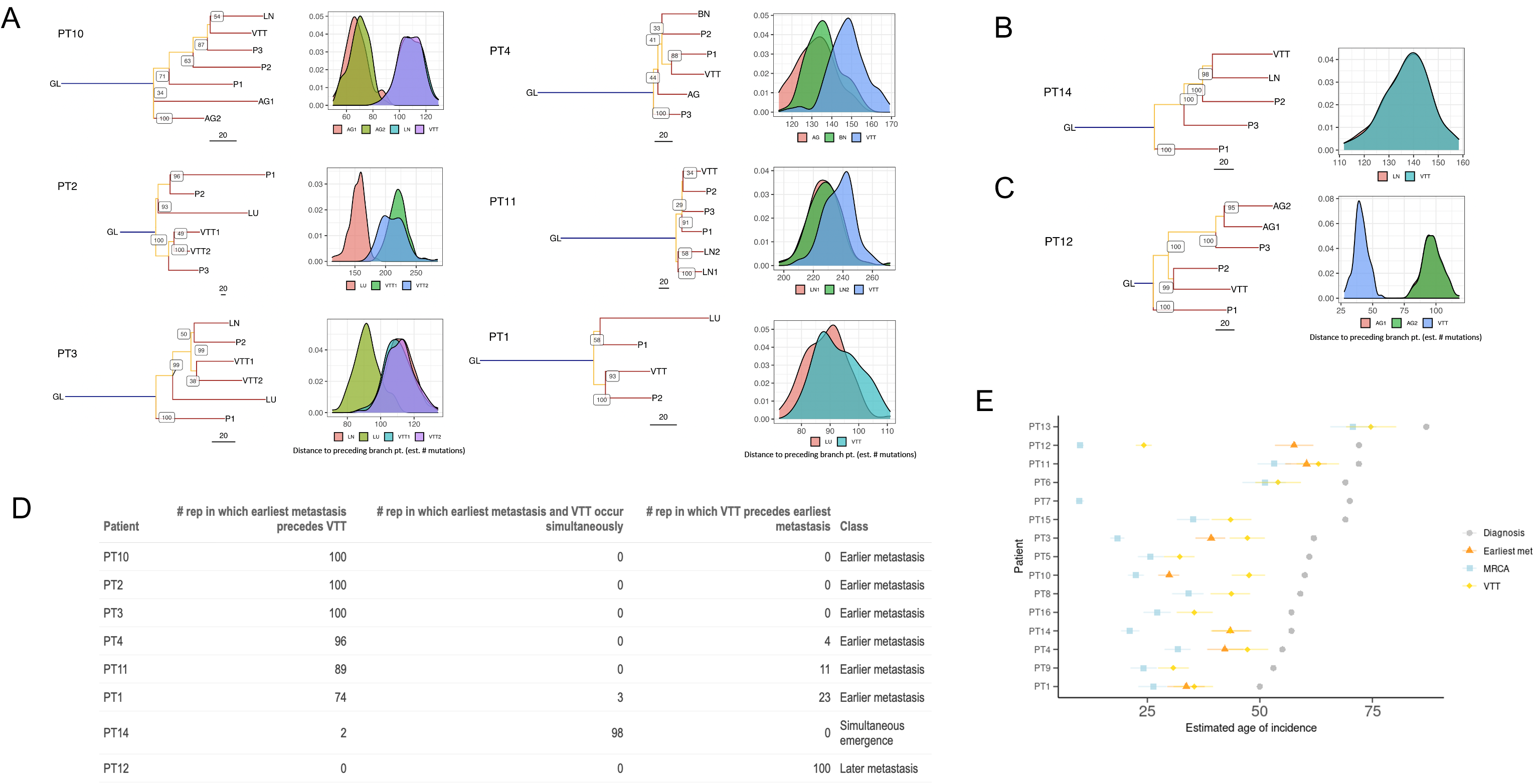
Phylogenetic and timing analyses of eight metastatic RCC tumors. (A-C) Mutational phylogenies and reconstruction robustness for eight RCC tumors with VTT and distant metastasis. Internal node labels on the phylogenetic trees display the number of bootstrap replicates supporting each node (i.e., the certainty out of 100 that the placement of a particular node is supported). Density plots illustrate the distribution of path lengths to the parent nodes of all non-primary regions derived from 100 bootstrap replicates of the tree building process. Region labels: GL = germline; P = primary tumor; VTT = venous tumor thrombus; AG = adrenal gland metastasis; LN = lymph node metastasis; LU = lung metastasis; BN = brain metastasis. (A) Patients with early emergence of metastases relative to VTT. (B) Patient with near-simultaneous emergence of VTT and the earliest assayed metastases, with both emerging from the same branch. (C) Patient with late emergence of metastases relative to VTT. (D) Table showing the number of bootstrap replicates in which phylogenetic tree reconstruction resulted in earlier, simultaneous, or later emergence of the earliest metastasis relative to the VTT. (E) Timing estimates generated with linear mixed effects models of the emergence of MRCA and earliest metastasizing clones, and the VTT clone, relative to the age at diagnosis. Error bars indicate 95% confidence intervals generated by parametric bootstrapping of the i.i.d errors.

The relative timing of a given phylogenetic node can be approximated by the sum of the internal branch lengths leading up to that node. However, a single “representative” tree does not fully capture the inherent uncertainty in these branch lengths. To make our nodal confidence analysis more quantitative, we developed a novel approach for quantifying total internal branch lengths that also captured the uncertainty in the phylogenetic reconstruction of the clonal ancestors of the VTT or of the metastasis. For each tumor, we quantified the total internal branch lengths across 100 bootstrap trees, and plotted the resulting empirical distribution of distances from the root node to branch points immediately preceding either VTT or metastasis. These root-node distance distributions then provide a measure of certainty regarding which region emerged first (Figure 3A-C, bottom panels). The frequency with which one branch-point precedes another in the bootstrap sampled trees provides an estimated p-value (Figure 3D). For example, in PT10 the AG1 and AG2 metastases showed clearly lower root-node distance distributions than the VTT and LN metastasis, suggesting that AG1 and AG2 emerged significantly earlier in mutational time. Similarly, in PT4 the AG and BN metastases showed consistently lower root-node distances than the VTT, with 96/100 bootstrap trees supporting the earlier emergence of the metastases (Figure 3A and D). Taken together, these results suggest that not only does the VTT typically not give rise to metastases, but many metastases actually emerge earlier than does the VTT.

This analysis revealed three categories of RCC tumors: (1) early emergence of at least one metastasis prior to VTT emergence, (2) late emergence of the earliest metastases after VTT emergence, and (3) similar timing of VTT and the earliest metastasis emergence due to their divergence from the same branch. Patients PT2, PT3, PT4, and PT10 fell clearly into the early metastasis class (p < 0.05, Figure 3A). Patient PT14 exhibited similar timing of relative emergence due to the VTT and metastasis sharing the same immediate ancestor (p < 0.05, Figure 3B). Patient PT12 fell clearly into the late metastasis class (p < 0.01, Figure 3C). Patients PT1 and PT11 were most consistent with the early metastasis class, but there was lower certainty (p ≅ 0.26 and p ≅ 0.11, respectively) in distinguishing the root-node distance distributions for VTT and metastasis owing to the small numbers of mutations distinguishing their immediate ancestors (Figure 3D). We note that in three patients (PT2, PT10, PT11), the earliest metastatic clones emerged prior to diversification of the earliest primary tumor clones.

Estimating the absolute timing of metastatic clone emergence could have important implications for clinical care of RCC. Having established the relatively early emergence of metastatic clones relative to VTT clones, we sought to determine the age of the patients at which metastases emerged, as well as the number of years between metastasis emergence and clinical diagnosis. We adapted the modeling approach of Mitchell et al (16) by generalizing it to encompass any node on the tumor phylogeny. This allowed us to infer the absolute timing of metastatic subclone emergence relative to the root (most recent common ancestor, or “MRCA”), to the VTT subclone, and to age at diagnosis. Using the average of the root-node branch lengths from each patient’s bootstrap sampled trees, we predicted the patient ages at which the earliest metastasis and VTT branching events occurred (Figure 3E). This timing analysis suggested that metastases emerged 11.5-30.3 years prior to diagnosis (Table S5) in the seven assessed patients (PT2 was excluded due to the *MLH1* mutation leading to violation of model assumptions). This analysis largely recapitulated the observations from the phylogenetic trees, with the early-metastasis patients also exhibiting earlier absolute estimates of metastasis emergence as compared to VTT emergence. Three of these patients appeared to show nearly concomitant emergence of the VTT and metastasis clonal ancestors (PT1, PT4, PT11), and the other two patients had significant, multi-year lags between the emergence of the VTT and metastasis clonal ancestors (PT3, PT10). In PT12, the sole patient in whom the emergence of the VTT preceded the earliest metastasis, the VTT clonal ancestor emerged a striking 33.4 years prior to the emergence of the adrenal gland metastases.

### TNF-α signaling is associated with VTT emergence, and MTOR signaling is associated with metastasis

To explore whether distinct molecular programs govern the outgrowth of VTT from primary tumor, as well as metastatic clone outgrowth, we performed paired differential gene expression analysis on 12 patients (RNA-seq was not available for PT1, PT2, PT3, and PT4). We first compared primary tumor regions to VTT regions while accounting for patient-specific variability in gene expression. We identified 30 genes as significantly upregulated in VTT relative to primary tumor, and 12 as significantly downregulated (Figure 4A and B).

**Figure 4.**
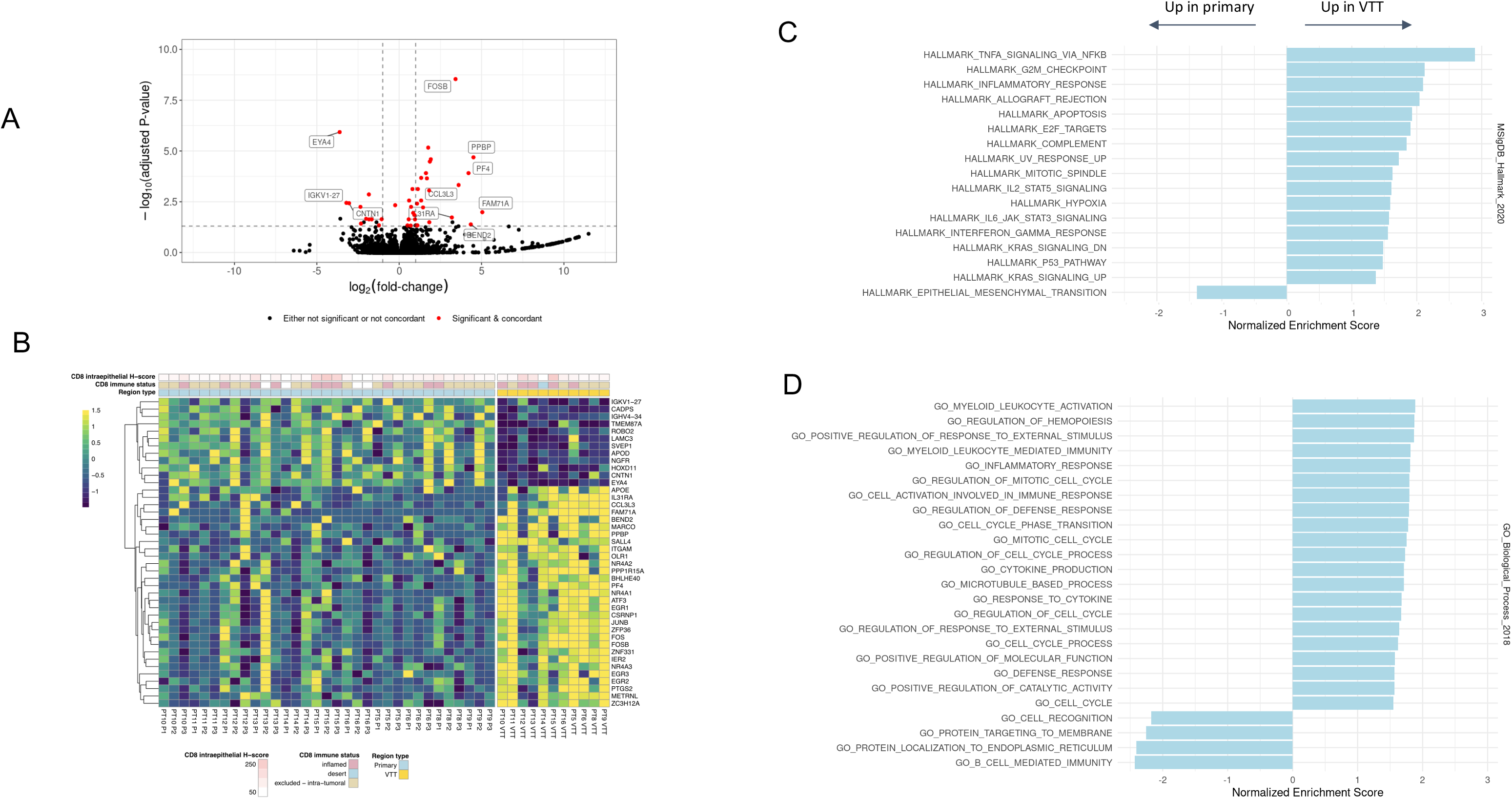
Differential gene expression analysis of VTT versus primary tumor regions. (A) Volcano plot showing log-fold changes and adjusted p-values for differentially expressed genes. Genes highlighted in red were both significantly different and exhibited concordance in the directionality of their changes across regions and patients. A few non-concordant outliers have been removed for clarity of plotting. (B) Heatmap showing expression values of differentially expressed genes between primary and VTT regions. Expression values are normalized per gene and per patient. (C) Significantly enriched or depleted Hallmark pathways in VTT relative to primary regions as determined by GSEA. (D) Significantly enriched GO Biological Process terms enriched or depleted in VTT relative to primary regions as determined by GSEA. Significance is defined as adjusted p-value <= 0.05 (Benjamini-Hochberg).

Gene set overlap analysis of the differentially expressed genes using Enrichr (33) revealed striking overlap with the TNF*α* signaling Hallmark pathway (Figure S6A). Out of the 42 differentially expressed genes, 17 are associated with TNF*α* signaling (adjusted p-value < 10^-6^). Gene set enrichment analysis (GSEA) with the full set of detected genes (see Methods) revealed a number of significantly upregulated Hallmark pathways (46), including inflammatory response (Figure 4C). Intriguingly, the sole significantly downregulated pathway was epithelial to mesenchymal transition (EMT). GSEA further revealed a number of significantly enriched gene ontology biological process (47) (GOBP) terms (Figure 4D). The upregulated GOBP terms were concordant with the observed enrichment of the Hallmark inflammatory response pathway, and included many immune-related processes, as well as some related to the cell cycle. An overlap of our differentially expressed gene list with genes identified by a previous study (4) resulted in three genes in common: *MARCO*, *PF4*, and *PPBP*. These genes as well as others that are upregulated in VTT (e.g., *APOE*) are associated with activated macrophages and platelets.

To address the possibility that these differences in gene expression were driven by specific cell types, we performed histology analysis and gene expression deconvolution analysis. The histology analysis confirmed that significant neutrophilic infiltrate was not detected in VTT or primary tumors, and IHC analysis suggested no increase in CD8 immune infiltration of VTT as compared to the primary tumor (Figure 4B). We then used xCell (48) to assess the enrichment of gene expression signatures associated with 67 immune and stromal cell types. We normalized the scores within each cell type and patient, and compared them across tumor region types (Figure S7). No cell type was significantly different across regions after correction for multiple testing. Together, these results suggest that TNF*α* signaling and inflammation-like changes in genes expression are associated with outgrowth of VTT from primary tumor, but the TNF*α* likely does not arise from lymphocytic infiltration of tumor.

We next compared primary tumor regions to metastases, and identified 19 genes as significantly downregulated in the metastases relative to the primary tumors, and five genes as upregulated (Figure 5A and B). Notably, *MTOR* was consistently upregulated in metastases relative to primary tumor regions. Although no terms were enriched in these 24 genes alone, GSEA with the full set of detected genes revealed an upregulation of MTORC1 signaling in metastases, along with *MYC* targets and oxidative phosphorylation (Figure 5B). These data are consistent with previous observations that mutations that increase mTOR activity are associated with metastatic RCC (49).

**Figure 5.**
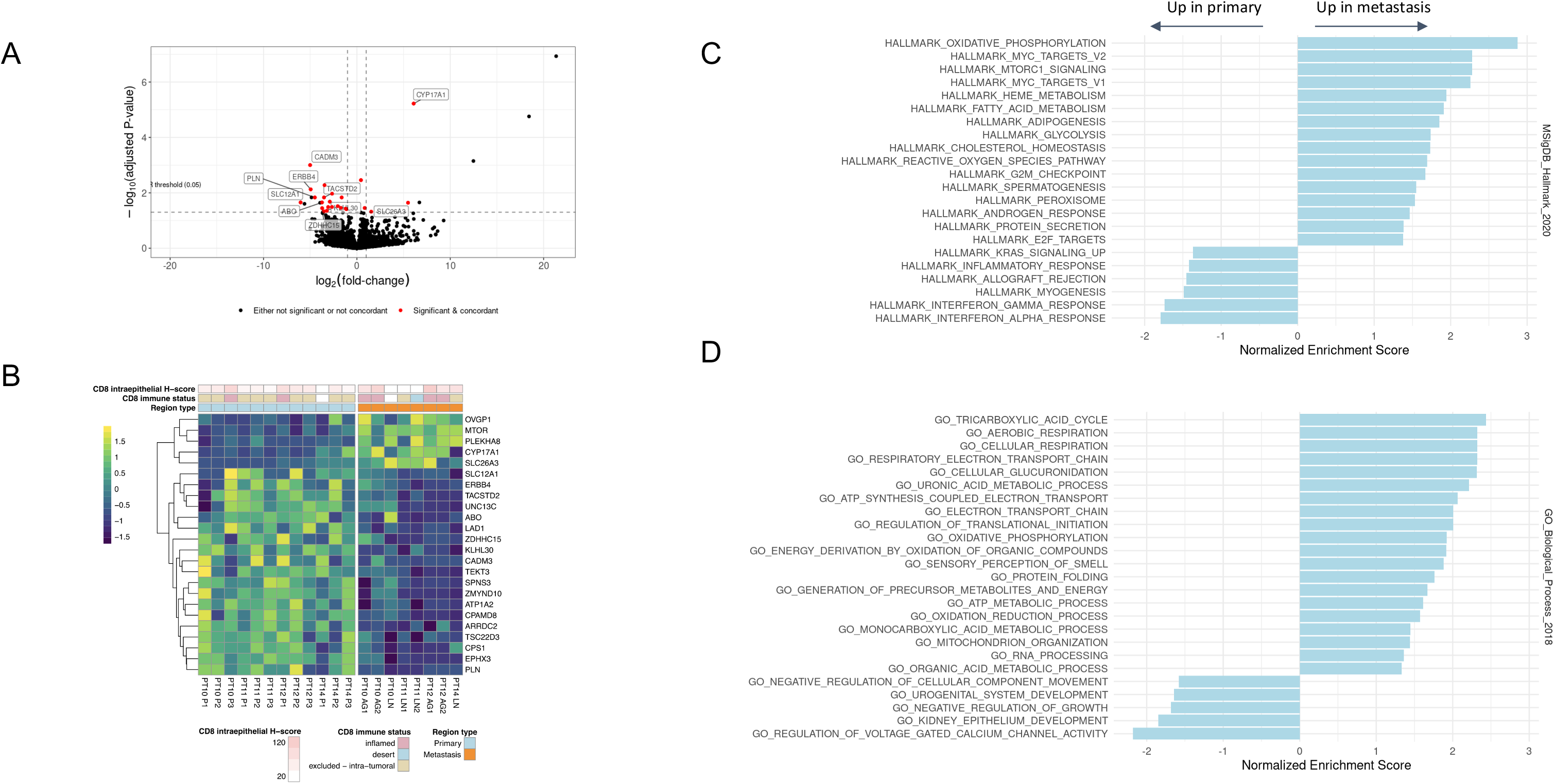
Differential gene expression analysis of metastases versus primary tumor regions. (A) Volcano plot showing log-fold changes and adjusted p-values for differentially expressed genes. (B) Heatmap showing expression values of differentially expressed genes between primary and metastatic regions. Expression values are normalized per gene and per patient. (C) Significantly enriched or depleted Hallmark pathways in metastases relative to primary regions as determined by GSEA. (D) Significantly enriched GO Biological Process terms enriched or depleted in metastases relative to primary regions as determined by GSEA. Significance is defined as adjusted p-value <= 0.05 (Benjamini-Hochberg).

Finally, we compared gene expression in the metastases to that in VTT, and identified four genes as significantly upregulated in the metastases relative to VTT, and five genes as downregulated (Figure S8A). This analysis was restricted to the four patients for which we had both metastatic samples and RNA-seq data, and so may be under-powered.

## Discussion

In this study we analyzed 16 RCC patients by MR-seq to characterize the genomic and evolutionary landscape of tumors with VTT. The unique nature of these cases--with a distinct lesion extending from the primary tumor and in some cases with separate metastases--also enabled us to investigate how and when new lesions arise from the primary tumor. Although only eight patients had both VTT and metastasis, our study surveyed a diverse variety of metastatic sites across these cases. Among metastatic cases, genomic alterations were generally consistent with advanced disease state. Given that RCC is known for frequent subclonal driver mutations, it was somewhat surprising that most tumors tended to share the more prevalent driver mutations across all regions (Figure 1A). Interestingly, the six tumors with higher regional heterogeneity among driver mutations also tended to have parallel mutations in separate tumor regions (Figure 1B). This may point to early RCC clones tending to grow in “fits and starts”, and only when a critical set of hallmark pathways become altered can full-blown tumor lesions emerge. We found that while 3/16 cases exhibit patterns consistent with the seven evolutionary subtypes proposed by Turajlic et al (7), others appear to follow distinct evolutionary modes (Table 1). Notably, 37% of the TRACERx RCC cases were also considered unclassifiable. We did not identify candidates for recurrently mutated genes restricted to the VTT, suggesting that tumor-intrinsic epigenetic factors or tumor-extrinsic factors may lead to VTT formation.

As has previously been reported, we observed extensive, early branching evolution in the ccRCC cases (Figure 3A-C and Figure S4). The two pRCC cases exhibited a greater degree of truncality (and later branching) than the metastatic ccRCC cases. At this time, too few pRCC cases have been analyzed by MR-seq to generalize this observation. Despite the genetic heterogeneity, mutational signatures were relatively concordant across regions, with occasional exceptions such as region P3 from patient PT12, which was the only region to exhibit substantial contributions from signature S5. We observed signature S6, which is associated with mismatch repair deficiency, consistently across most patients. The presence of this signature, albeit at low levels in most cases, is seemingly at odds with the relatively low mutation burden broadly observed in RCC (29). Given that many RCC tumors respond to immunotherapy agents, future studies should explore whether the level of S6 represents a potential biomarker. We also observe signature S3, or “BRCA-ness”, in a substantial proportion of patients, consistent with previous observations (5). Genes required for homologous recombination and mismatch repair are downregulated by *VHL* inactivation, presumably via the resulting stabilization of HIF-1ʱ (50). The related presumed deficiency in double-strand break repair has generated recent interest in the use of PARP inhibitors to treat RCC (51), and the level of signature S3 may also represent a potential biomarker.

Our phylogenetic analysis of eight metastatic cases with VTT ruled out the possibility of direct hematogenous seeding of metastases from the VTT, with the possible exception of PT14, in which the VTT and LN metastasis share a clonal ancestor. Among these eight cases, we identified three primary modes of interplay between VTT and metastasis: early metastasis, late metastasis, and concomitant metastasis arising from the same parent clone. The latter two modes are consistent with what was described for the six metastatic cases with VTT in the TRACERx cohort. However, most tumors in our cohort (6/8) displayed a distinct early emergence of metastases relative to VTT. Indeed, in PT10 the two adrenal gland metastases were the first to emerge from the root node. A caveat of this analysis is that we are reliant on the use of the parent node in the phylogenetic tree as a proxy for the emergence of each tumor region or lesion. We cannot be certain that VTT or metastatic competence was attained at the time of this branching event, as it could have occurred later via subsequent mutations or epigenetic processes. Our findings that VTT development may be separate from the development of metastatic disease is consistent with clinical experience where aggressive surgical intervention despite VTT can result in long-term survival. Moreover, this apparent independence is consistent with the fact many patients with VTT present without metastatic disease, and the fact that probability of metastatic presentation is proportional to primary tumor size, while probability of VTT exhibition is not (52).

Our timing analysis enabled us to estimate absolute timing of metastatic clone emergence, as well as VTT emergence. This approach is contingent on the assumption that the tumor mutation rate is not substantially different from the background somatic mutation rate, and that mutation rates do not vary substantially across tumor regions. In our patients, the presence of a number of non-age associated mutation signatures suggests that the former assumption may be violated. Indeed, two of the mutational signatures we observed, S3 and S6, have been associated with higher SNV mutation rates (53,54). However, upon fitting the model, the patient-specific slope estimates were relatively similar to one-another, with the exception of PT2 (Figure S5). The truncal inactivating *MLH1* mutation likely led to an increased mutation rate in this tumor, and we therefore excluded PT2 from this analysis.

The results of the timing analysis suggested that in all patients the earliest metastases (i.e., the clones that ultimately constituted the earliest metastases) emerged well before diagnosis--as long as three decades in one patient. This early metastatic clone emergence is consistent with recent findings that metastases in colorectal, breast, and lung cancer are disseminated early relative to time of diagnosis (13,14). A caveat is that our timing analysis provides an estimate of the emergence time for a metastatic subclone--it does not establish when that subclone seeded a distal site. However, given that a large number of RCC cases present with metastases on diagnosis, early detection studies are needed to examine whether RCC metastases are formed well before the disease is clinically evident or detected. A better understanding of the patterns and timelines of RCC evolution has relevance not only for small renal masses but also for more advanced disease. The increased application of percutaneous tumor biopsy prior to treatment may allow personalized management based on molecular characterization of the cancer.

Differential expression analysis revealed 42 genes that were consistently differentially expressed between primary tumor and VTT. VTT regions showed upregulation of TNF*α* signaling (17 of 42 genes) and related inflammatory processes relative to primary tumor regions. There was no evidence from IHC analysis or immune-cell gene expression signatures that these differences were due to lymphocytic infiltration of VTT. It is possible that the source of TNF*α* in these RCC tumors was myeloid, stromal, or endothelial cells. RCC tumors are known to have an angiogenic phenotype and substantial myeloid involvement, leading us to speculate that intratumoral vascular endothelial cells and/or activated macrophages produce TNF*α*, and potentially other cytokines, leading the tumor to invade the renal vein.

The sole significantly depleted Hallmark pathway in the VTT identified by GSEA was epithelial-to-mesenchymal transition (EMT). This could indicate that the VTT is in general less competent to generate metastases than the primary tumor, rendering it a less frequent source of hematogenous metastases than might otherwise be expected. Although the genes we identified as differentially expressed between primary and VTT regions had minimal overlap (only three genes) with those from a previous study (4), the two analyses did identify similar pathways as upregulated (e.g., inflammatory response) and downregulated (e.g., EMT). A smaller number of genes were observed to be differentially expressed between primary and metastatic sites, but *MTOR* notably featured among them, with mTORC1 targets further featuring in GSEA results. *MTOR* signaling is thought to drive invasiveness in RCC and other cancers, and the use of mTORC1 inhibitors has seen some clinical success (49).

Taken together, our data reveal new dimensions of RCC VTT and metastasis biology. They highlight the need for earlier detection of RCC, and suggest that metastatic clones may emerge and remain largely dormant for years before becoming clinically detectable. They also suggest potential expression and mutational biomarkers of RCC patient prognosis and treatment response.

## Acknowledgments

We thank our colleagues at Genentech and at UCSF for helpful discussions.

## Supplementary case description

All 16 patients had VTT extending at least into the renal vein, and in nine it extended into the inferior vena cava. Seven of the 16 patients exhibited metastases on diagnosis (VTT+met), and metastases were later discovered in an additional four. The primary tumor regions and VTT samples were all derived from the initial VTT thrombectomy, which was performed shortly after diagnosis (in all but one case). Metastatic regions were sampled both from metastasectomies concomitant to the initial surgery, and from recurrences. In three of the six cases where recurrences were sampled, one or more non-surgical interventions preceded the sampling (PT1, PT4, PT11; see Figure S1, Table S1). Semi-quantitative histologic analysis of the 78 tumor regions revealed variable stromal abundance within tumors that was relatively consistent across tumor regions in any one patient. Very little intrastromal inflammatory response or desmoplasia was detected (Figure S2).

## Supplementary Methods

### AVENIO Millisect tissue harvest

AVENIO Millisect automated dissection for tumor enrichment was performed on all cases. Formalin Fixed Paraffin Embedded (FFPE) tissue blocks were serially sectioned with one section at 4µm, followed by 7 sections at 10µm, followed by 3 sections at 4µm, collected onto Superfrost Plus positively charged slides (Thermo Scientific, Runcorn, UK) and allowed to dry at room temperature overnight. Serial sections 1 and 9 (4µm) were baked at 60°C for 30 minutes and stained with Hematoxylin and Eosin (H&E) on an automated Leica Autostainer XL using a routine protocol. H&E stained slides were scanned on a NanoZoomer 2.0 HT whole slide imager (Hamamatsu, Bridgewater NJ) at 20X magnification. Scanned slide images were annotated by a pathologist for tumor regions of interest, percent tumor area necrosis (% necrosis/total tumor area) was captured and digital masks were created as a dissection reference.

Tissue sections were dissected using the reference mask image from serial section 1 to collect regions of interest using medium or large AVENIO Millisect milling tips (Roche Sequencing Solutions, Pleasanton, CA), collected with Molecular Grade Mineral Oil (Sigma-Aldrich, St. Louis, MO) as dissection fluid and dispensed into nuclease-free 1.5mL Eppendorf tubes. Dissections from slides 2 through 5 and 6 through 8 were centrifuged for 10 minutes at 20,000rpm to pellet tissue. Portions of mineral oil were removed from the tissue pellets. Pellets from slides 2 through 5 were pooled in a 1.5mL Eppendorf tube and held for DNA extraction and pellets from slides 6 through 8 were pooled in a separate 1.5mL Eppendorf tube and held for RNA extraction. Post AVENIO Millisect dissected tissue slides were baked at 60°C for 30 minutes and stained with Hematoxylin and Eosin (H&E) on an automated Leica Autostainer XL using routine protocols and scanned on a NanoZoomer 2.0 HT whole slide imager (Hamamatsu, Bridgewater NJ) at 20X magnification. DNA extraction was performed using the Qiagen AllPrep DNA/RNA tissue kit (Qiagen, Germantown, MD) at Q^2^ Solutions (Valencia, CA).

Tumor content ranged from 5 to 99% in analyzed tissue regions. Tumor enrichment was performed using AVENIO Millisect for semi-automated dissection, resulting in tumor input of 5.9-1439.81mm^2^ (Table S2) that excluded the majority of surrounding normal tissue and necrotic regions from capture and analysis.

### Estimating lesion emergence timing

In order to estimate absolute timing of lesion parent clone emergence, we adapted the linear mixed modeling approach of Mitchell et al (38), using the lme4 package (39). Patient age was modeled as a function of sample mutation counts as both a fixed and random slope, with intercept set to zero:

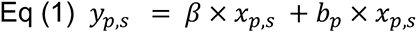

where *y_p,s_* is the age of patient *p* upon collection of sample *s*, β a fixed slope term, *x_p,s_* is the number of mutations detected in sample *s* from patient *p*, and *b_p_* is a random slope term for patient *p*.

Lesion parent clone emergence timing was estimated by predicting the patient age given the estimated number of mutations present at the branch point immediately upstream of the lesion on the phylogenetic trees. The number of mutations was estimated by calculating the sum of the branch lengths leading to that branch point in each bootstrapped tree, and then taking the mean over all the resulting sums. For example, the number of mutations at VTT emergence in PT4 was estimated by summing over the branch lengths leading to the node immediately upstream of the VTT (i.e. the branch point leading to both P1 and VTT in the representative tree in Figure 3A) in each of the 100 bootstrapped trees generated for that patient, and then calculating the mean of the resulting sums.

The timing of clone emergence was estimated for both the VTT (in all patients) and for the earliest metastasis (i.e. the metastasis with the fewest estimated mutations according to the above approach) in all patients from which a metastatic sample was collected. Confidence intervals were estimated using parametric bootstrapping of the model residuals with the *bootMer* function from the lme4 package.

## Supplementary Figure Legends

**Figure S1.**
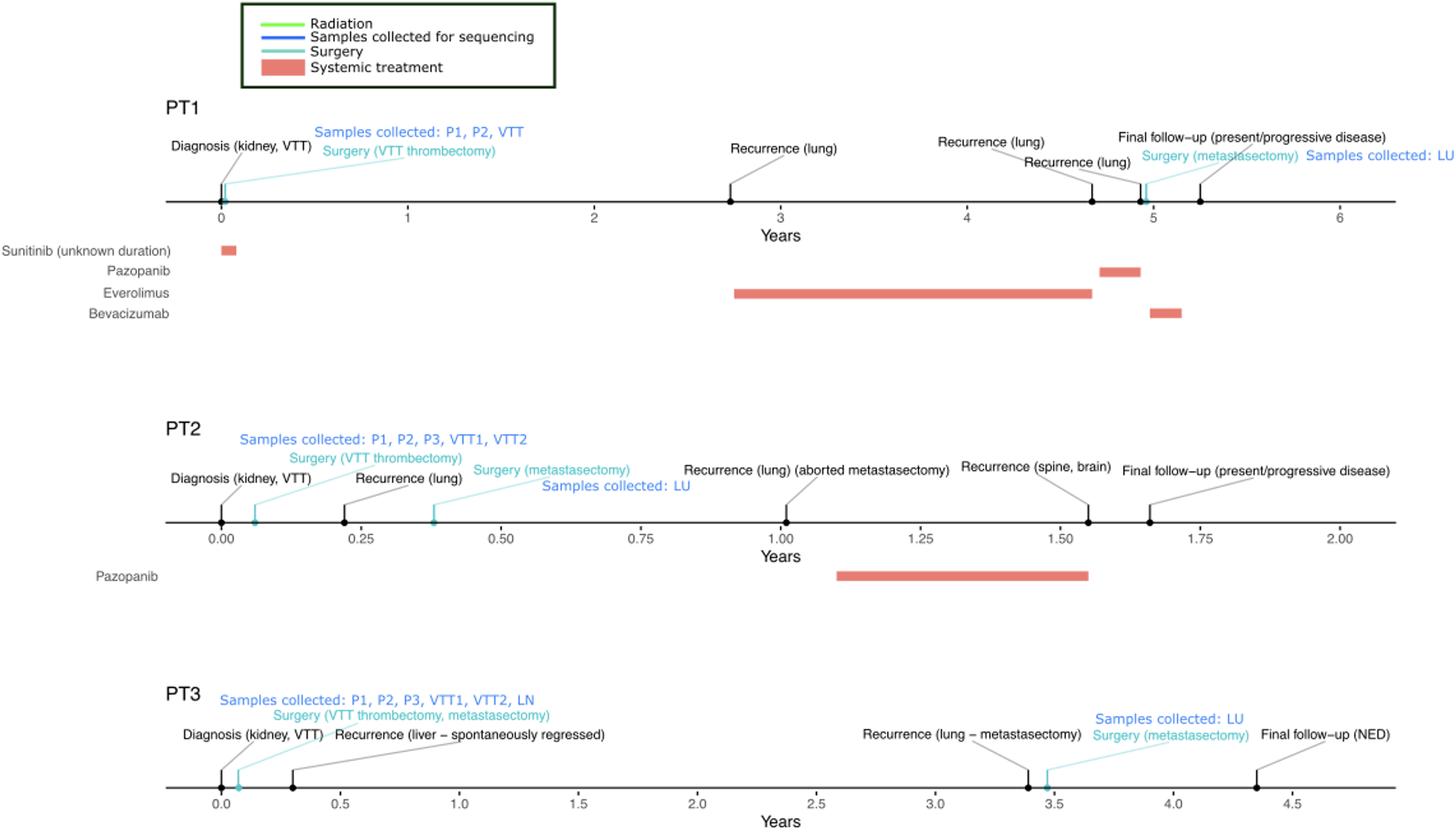
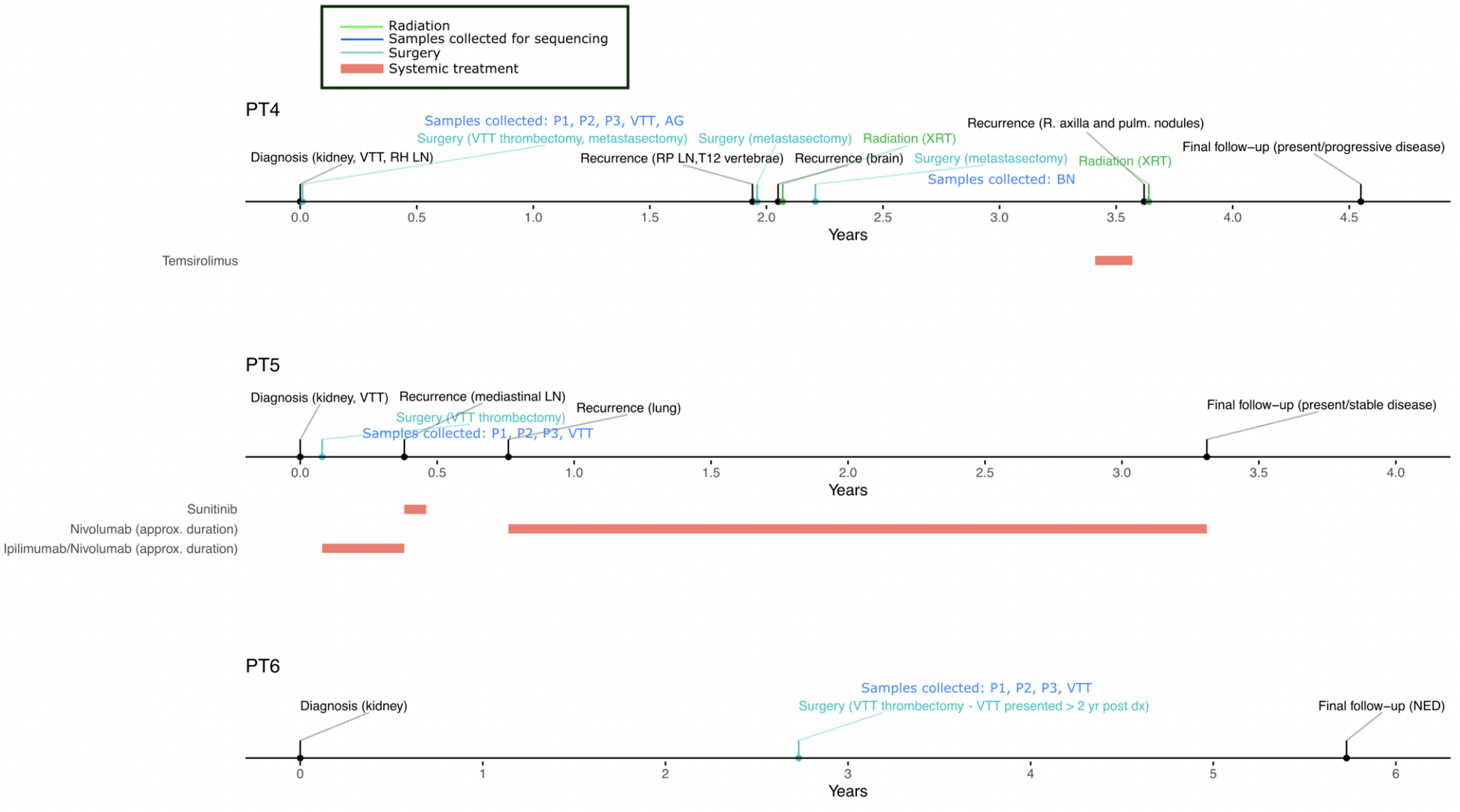
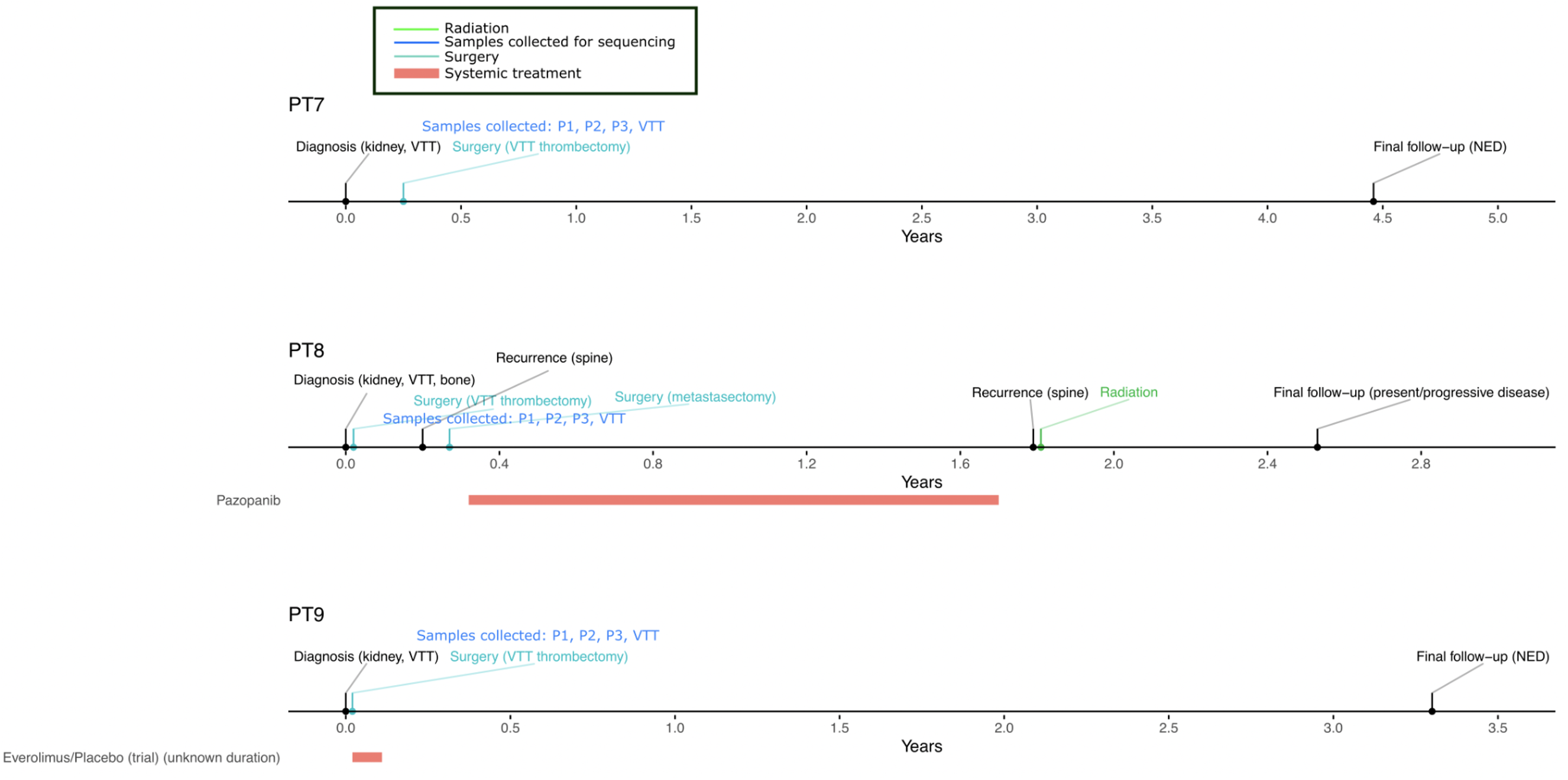
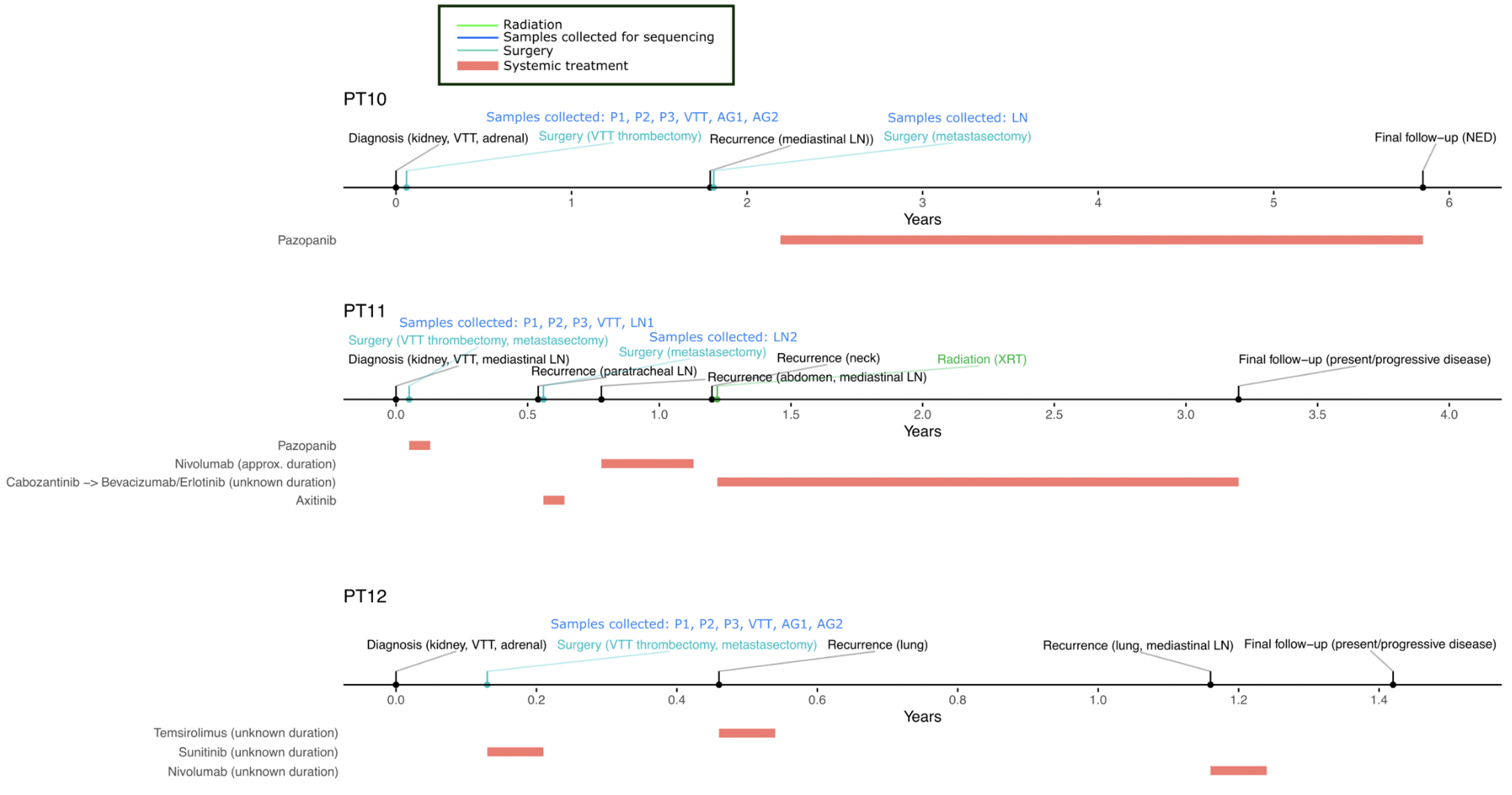
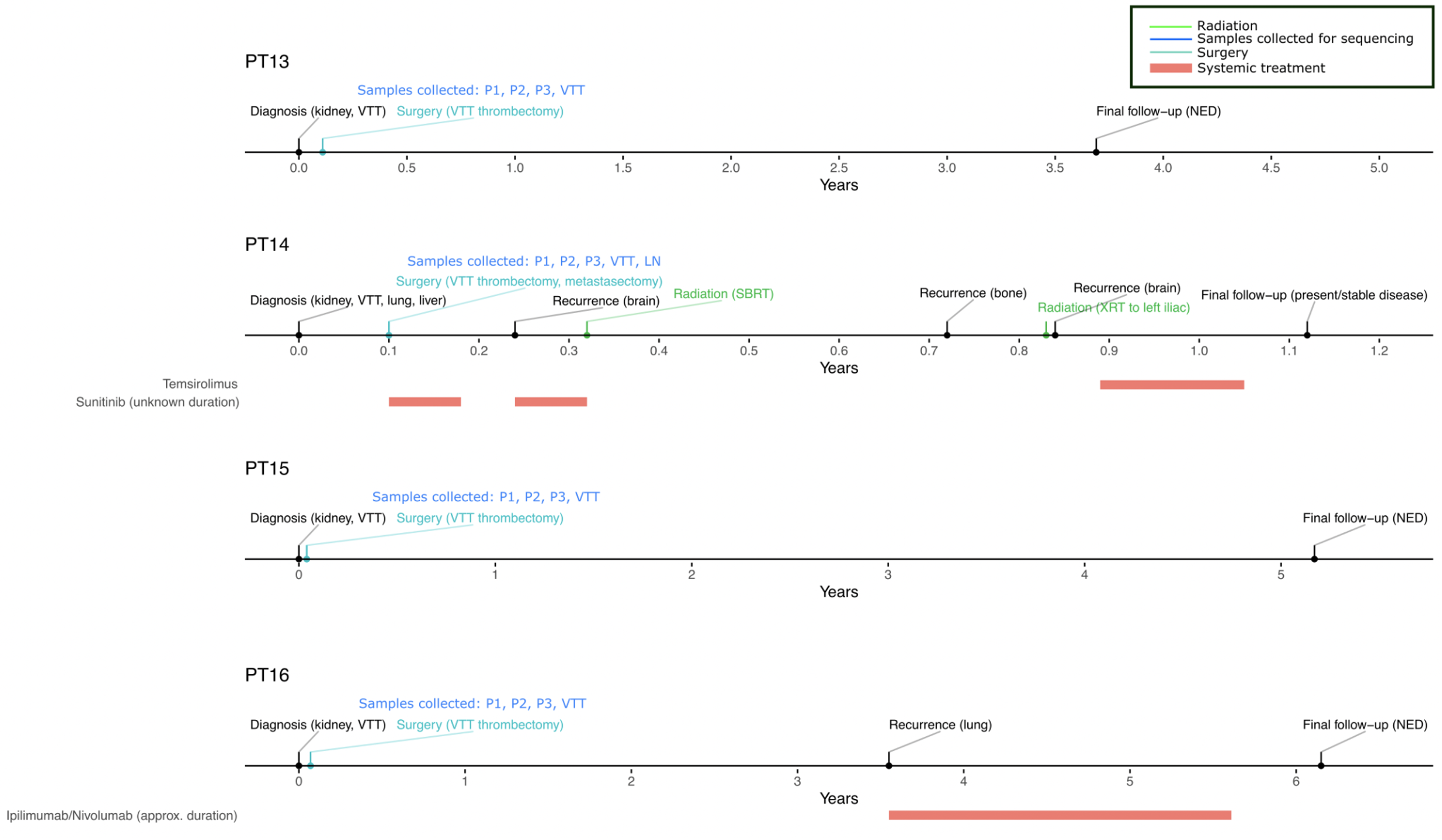
Timelines of patient diagnosis, surgeries, and other treatment.

**Figure S2:**
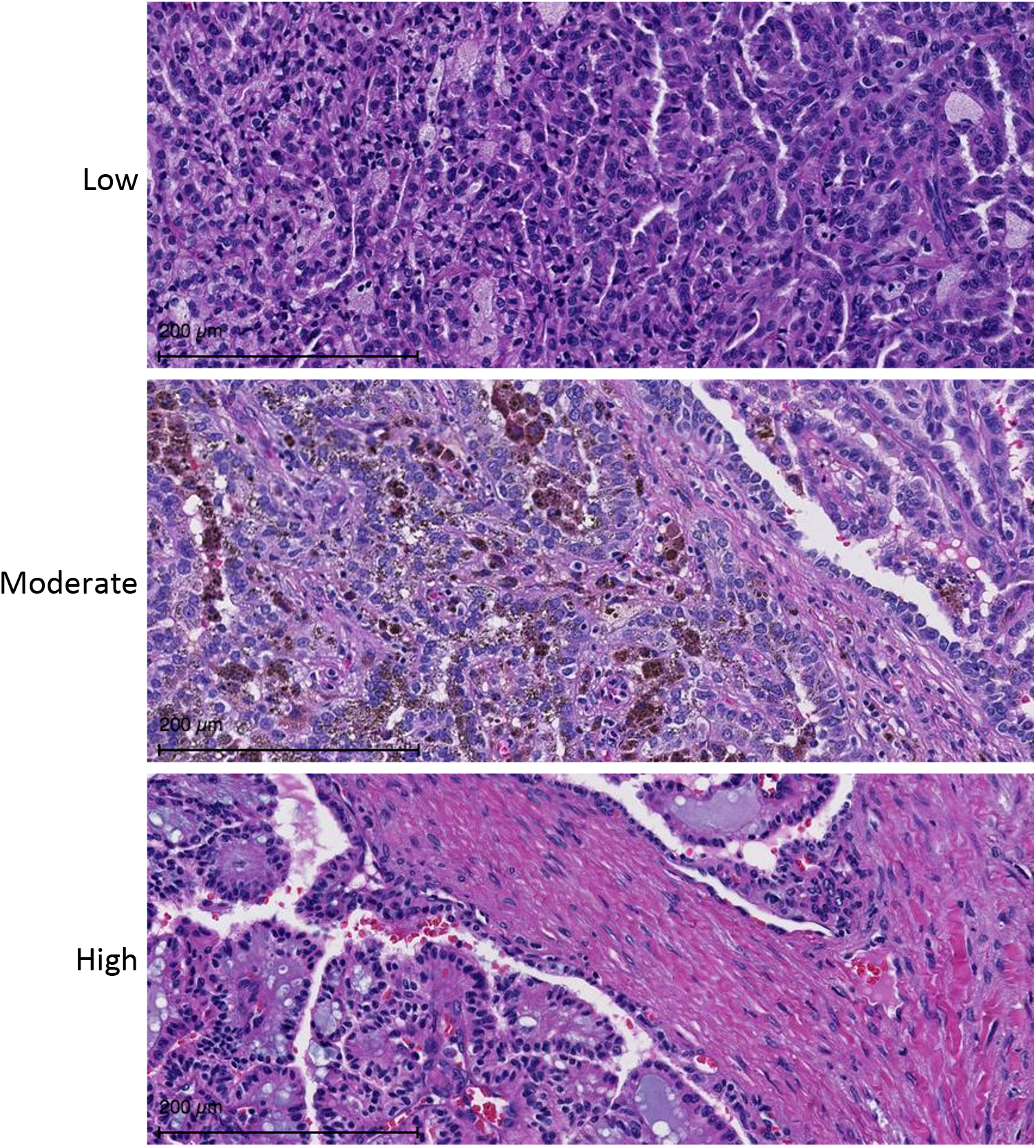
Semi-quantitative analysis of stromal abundance captured 3 levels (low, moderate or high) of stromal presence within tumors with little inflammatory response.

**Figure S3.**
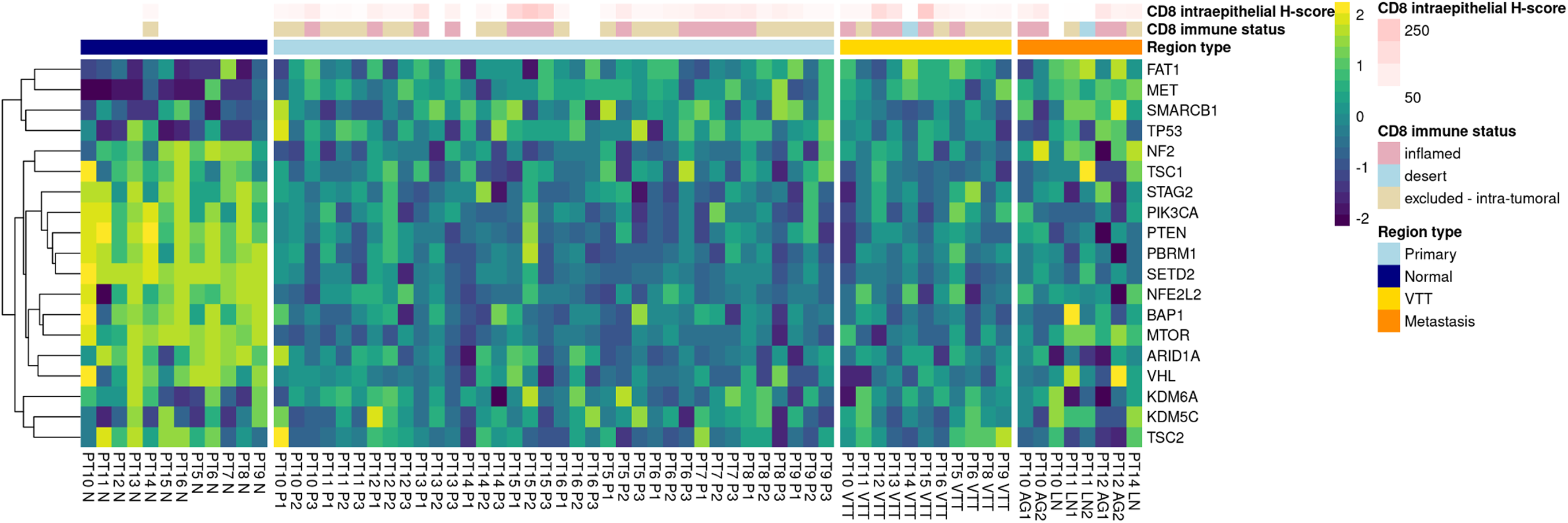
Expression status of putative driver genes across all regions of all patients, excepting CSMD3, which had no uniquely mapped counts in any region. Expression values were normalized by variance-stabilizing transformation, then turned into z-scores normalized within each gene and patient prior to visualization.

**Figure S4.**
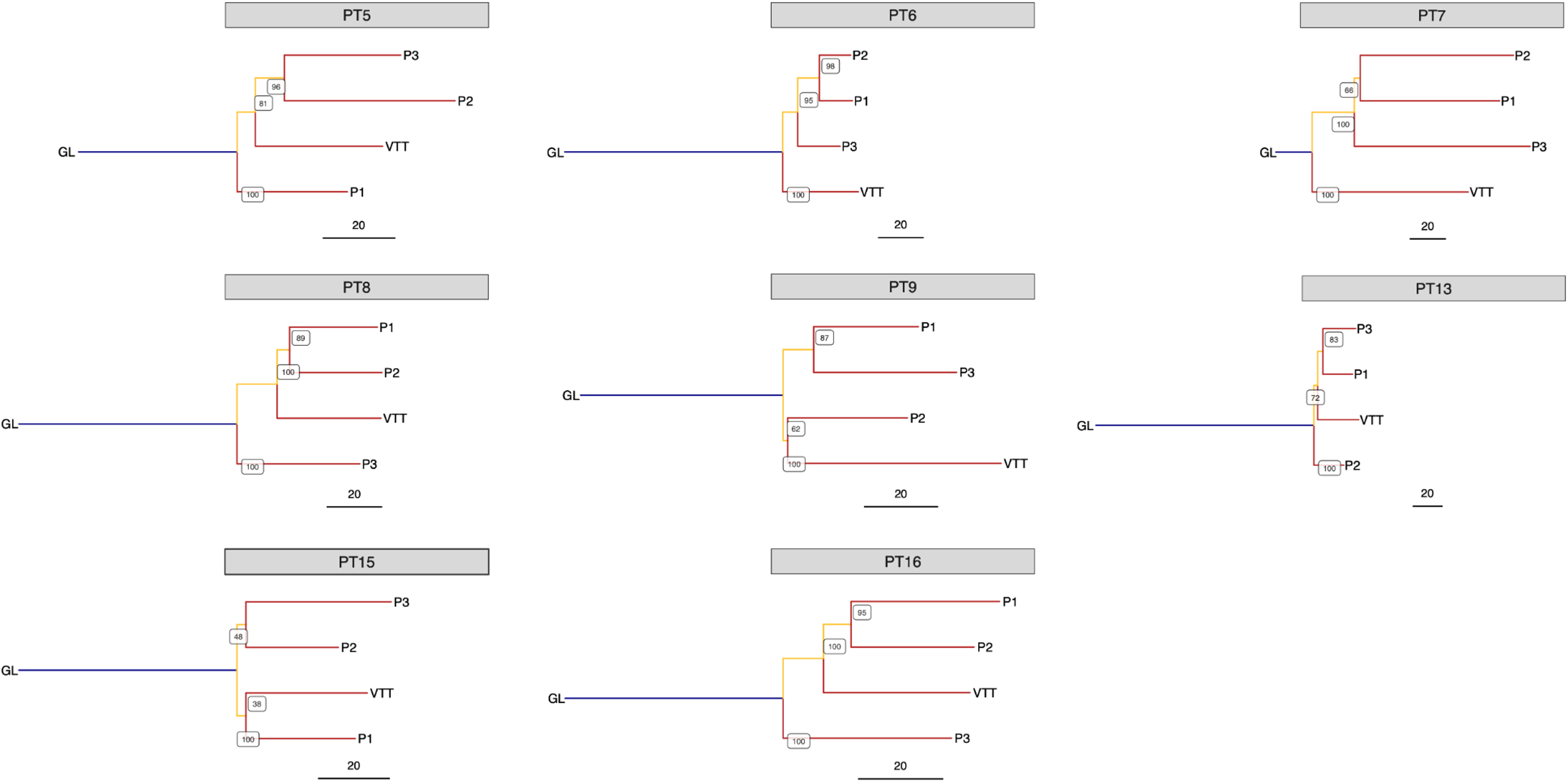
Phylogenetic analysis of eight RCC tumors, which are either non-metastatic or are missing metastasis samples. Internal node labels on the phylogenetic trees display the number of bootstrap replicates supporting each node (i.e., the certainty out of 100 that the placement of a particular node is supported). Region labels: GL = germline; P = primary tumor; VTT = venous tumor thrombus.

**Figure S5.**
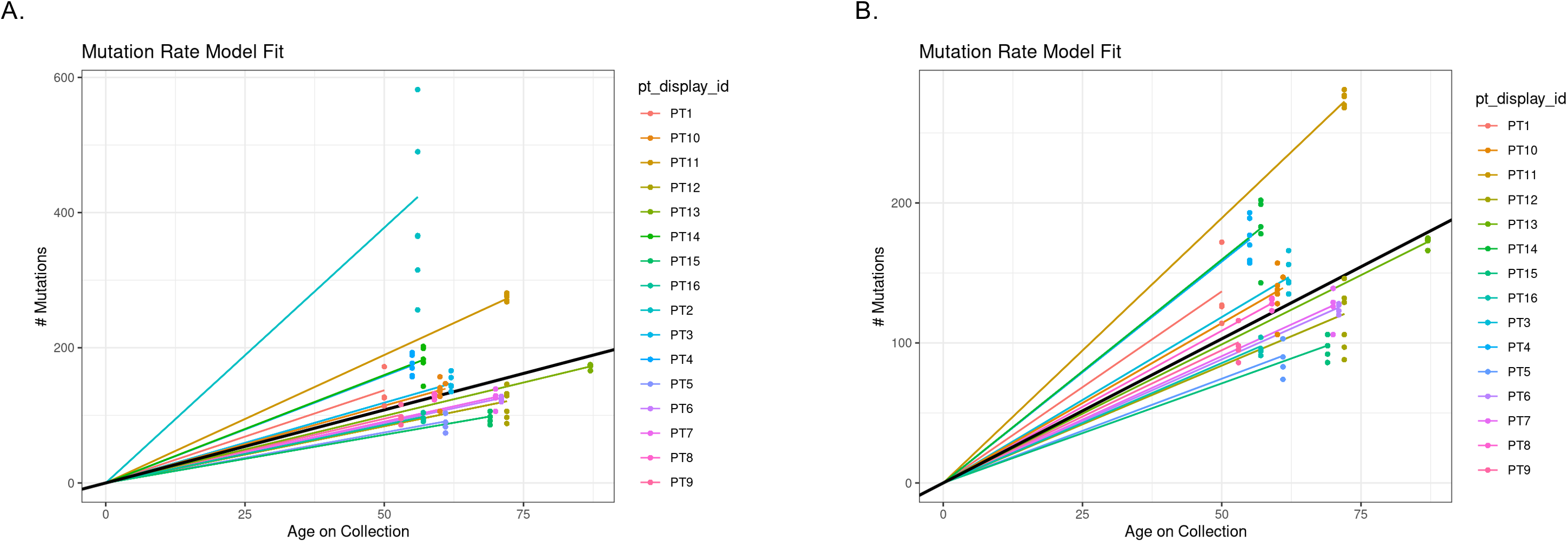
LME mutation rate model fits (A) with and (B) without PT2.

**Figure S6.**
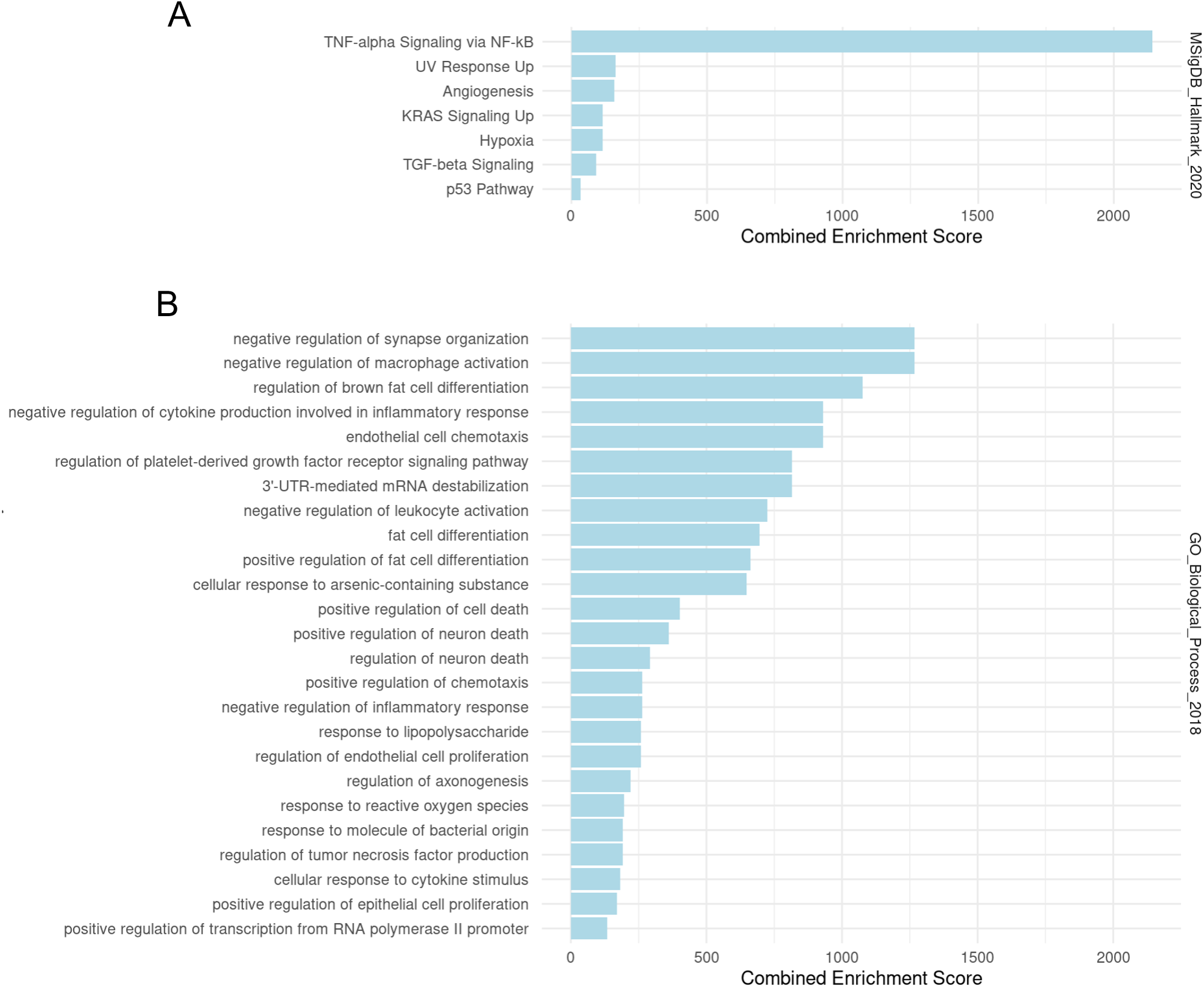
Enrichr set enrichment of differentially expressed genes between primary and VTT regions. (A) Significantly enriched terms from MSigDB Hallmark pathways. (B) Significantly enriched terms from GO Biological Processes. Significance is defined as adjusted P-value <= 0.05.

**Figure S7.**
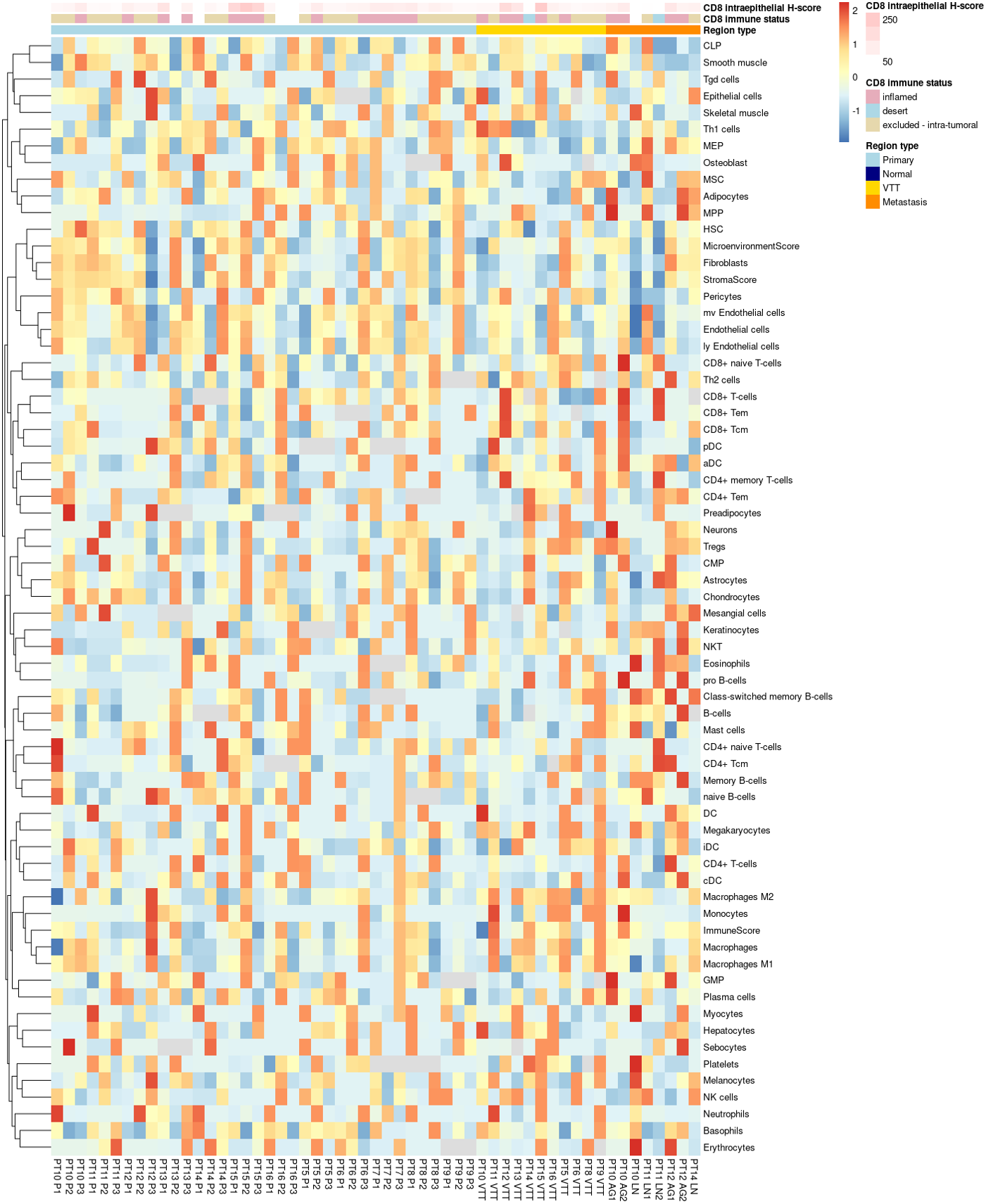
xCell enrichment scores for 67 cell types across all regions of all patients. Scores were normalized within each cell type and patient prior to visualization.

**Figure S8.**
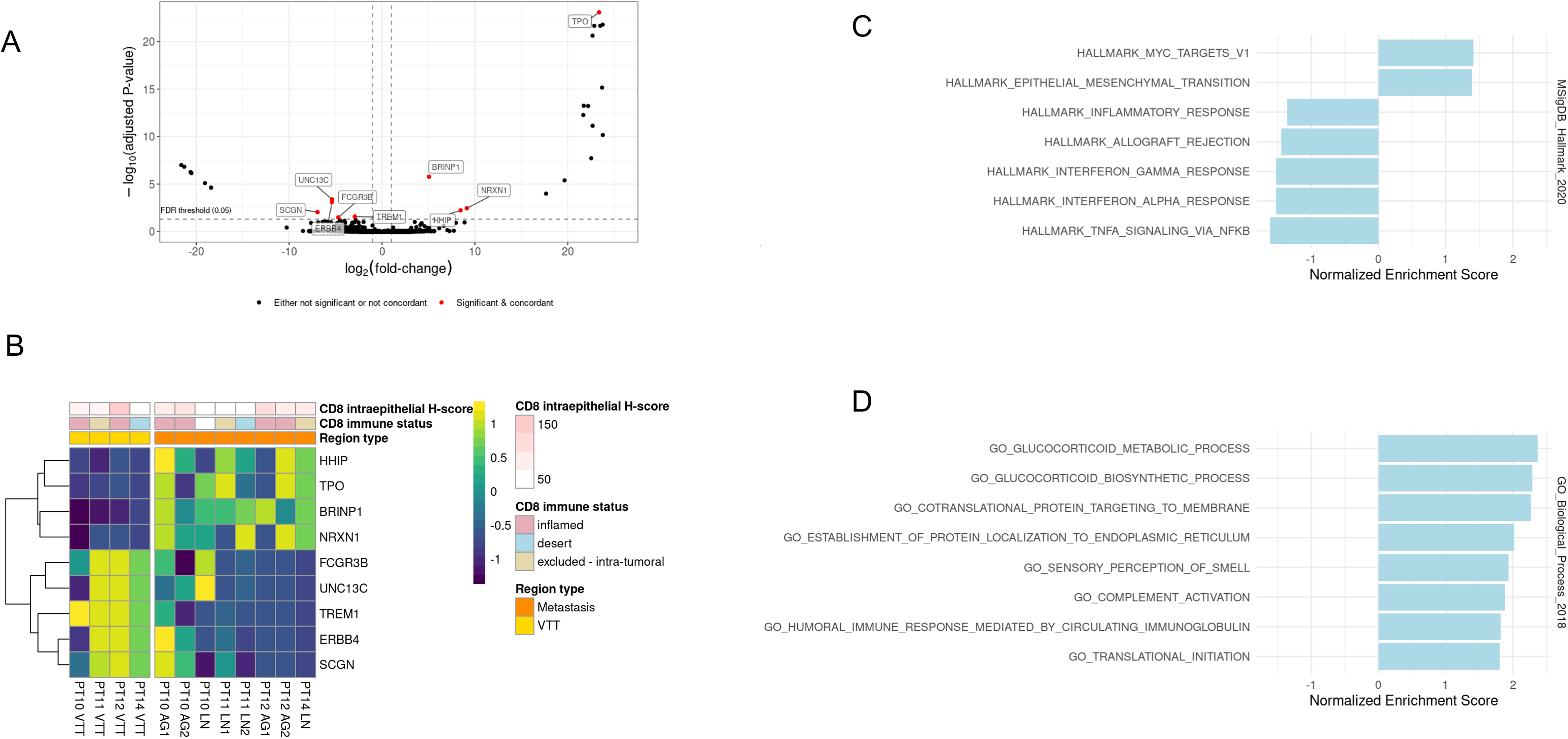
Differential expression between VTT and metastatic regions. (A) Volcano plot showing log-fold changes and adjusted P-values for differentially expressed genes. (B) Heatmap showing expression values of genes determined to be differential between VTT and metastatic regions. Expression values are normalized per gene and per patient. (C) Significantly enriched or depleted Hallmark pathways in metastases relative to VTT regions as determined by GSEA. (D) Significantly enriched GO Biological Process terms enriched or depleted in metastases relative to VTT regions as determined by GSEA. Significance is defined as adjusted p-value <= 0.05 (Benjamini-Hochberg).

**Table S1.**
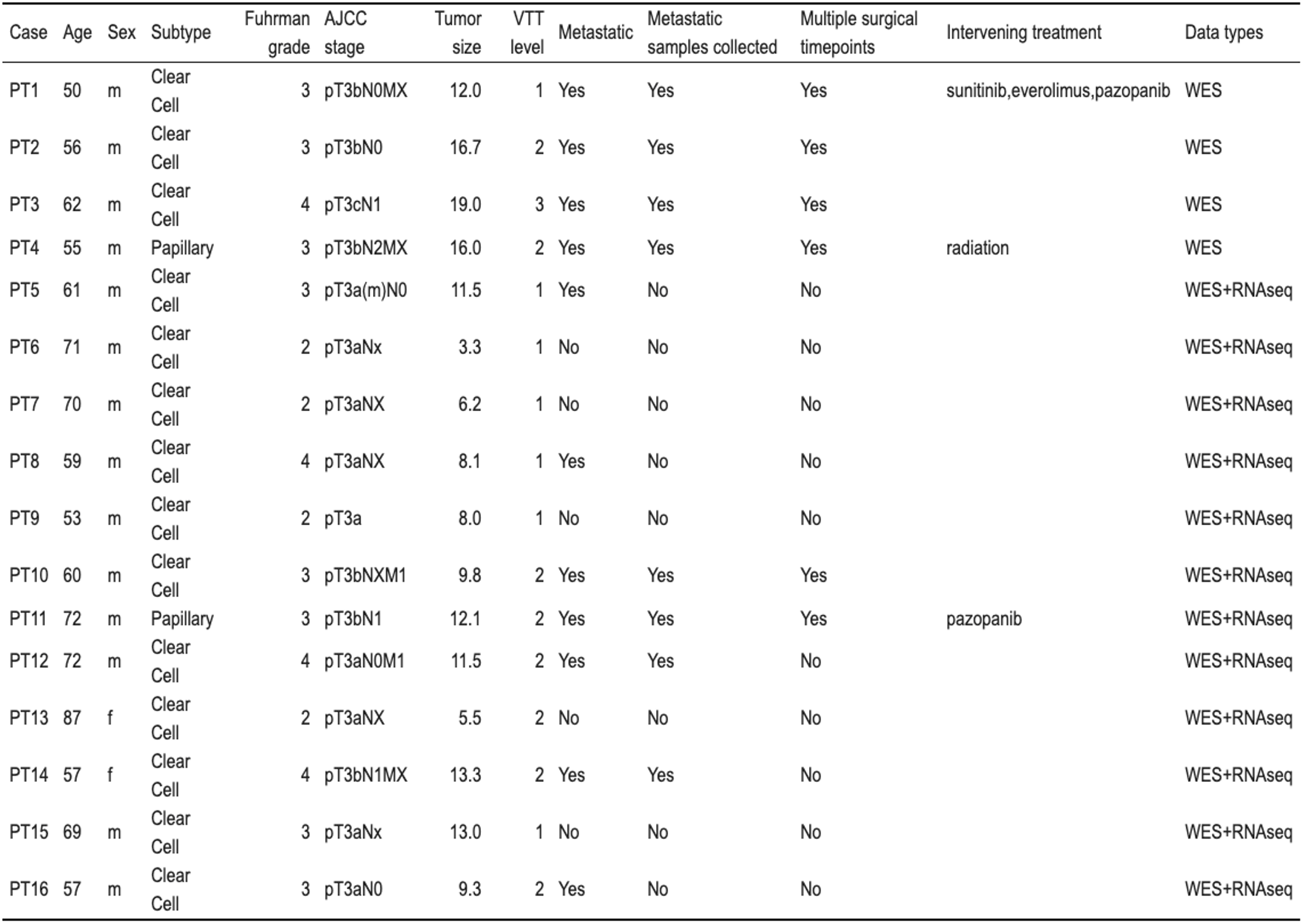
Extended characteristics of cases in this study.

**Table S2.**
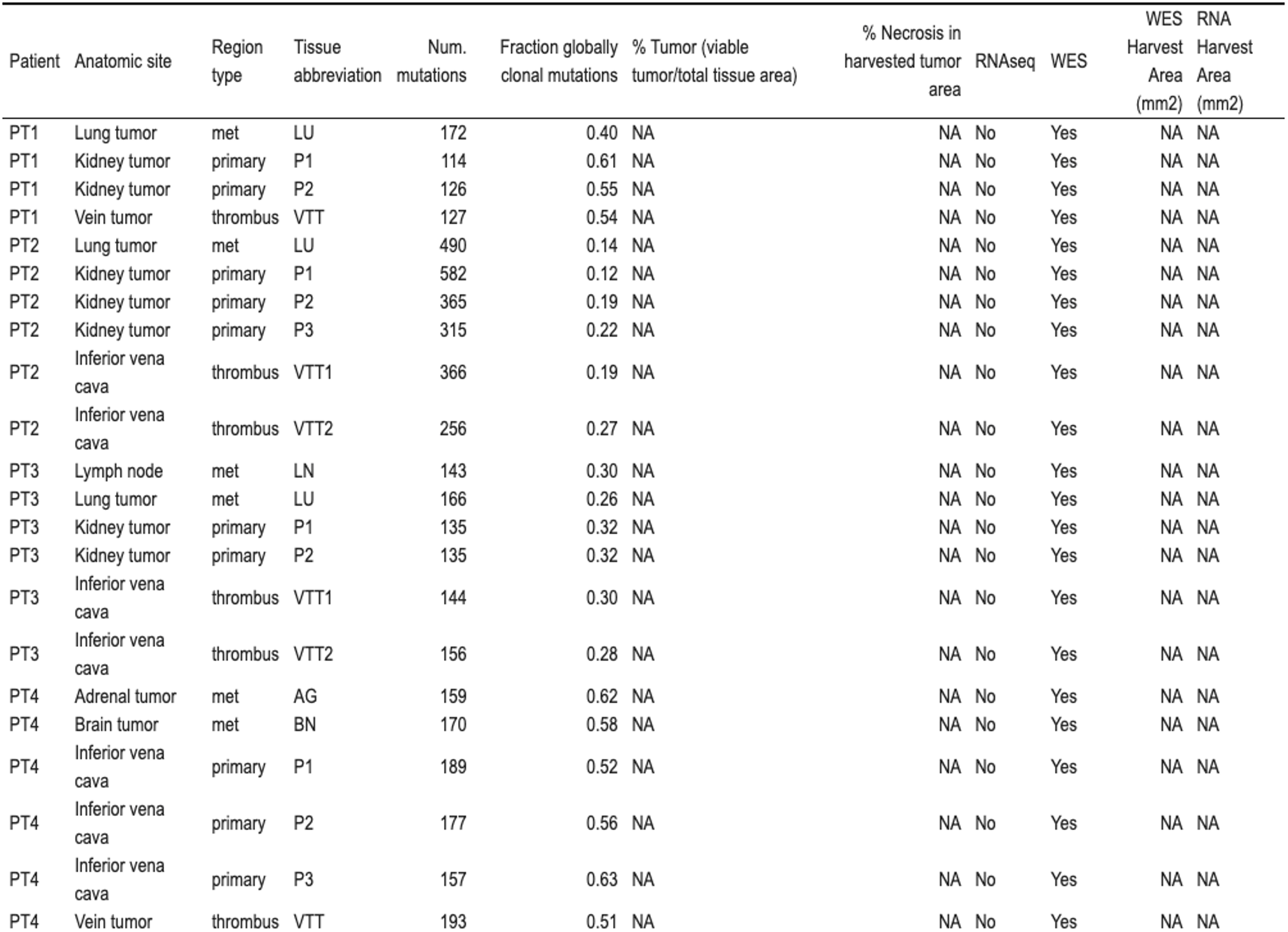
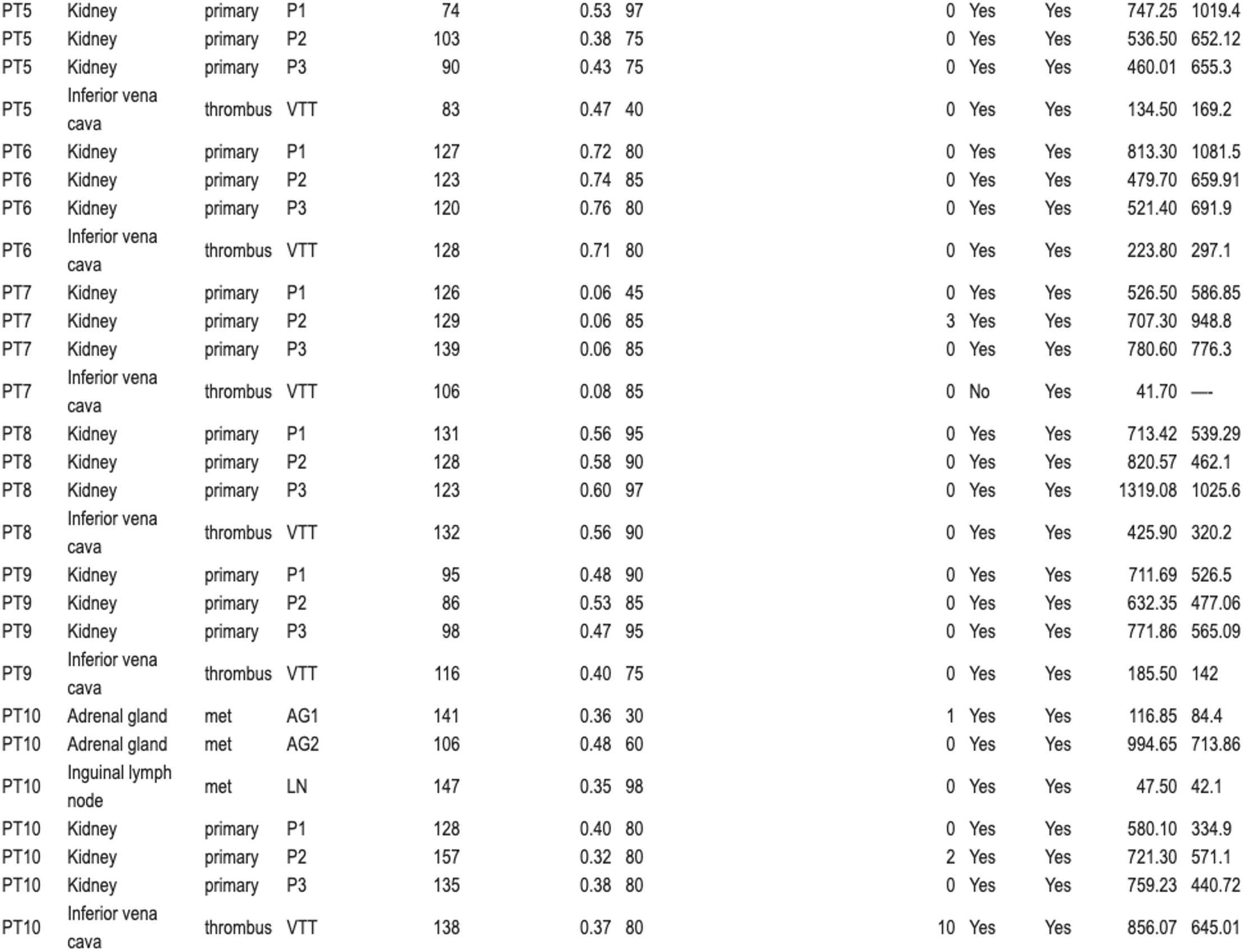
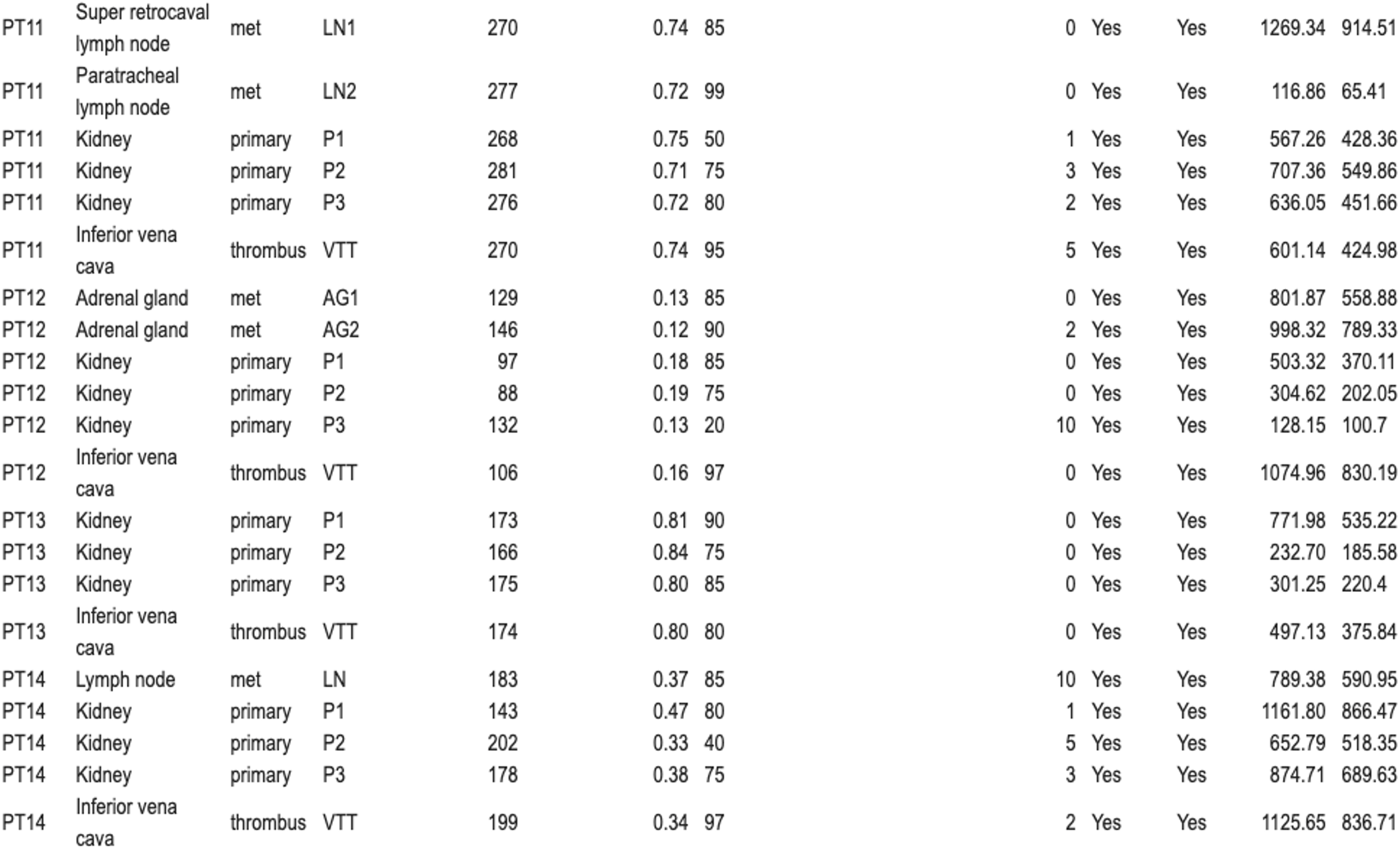
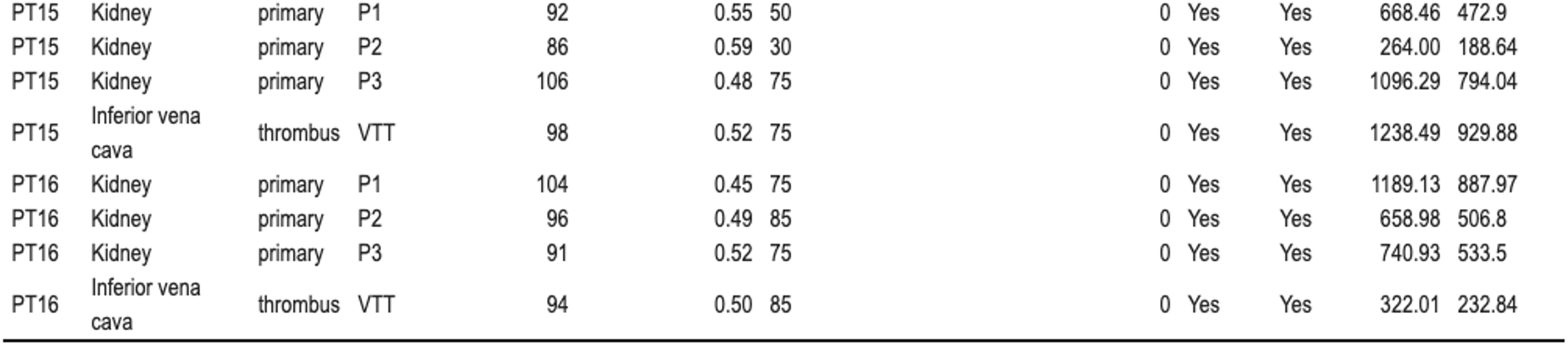
Characteristics of individual samples in this study.

**Table S3.** Mutation burden and across patients.

| Patient | Total unique mutations | Globally clonal mutations | Mean mutations/sample | Mean mutations/sample/Mb | Mean fraction globally clonal |
| --- | --- | --- | --- | --- | --- |
| PT1 | 249 | 69 | 130 ± 26 | 2.7 ± 0.51 | 0.52 ± 0.087 |
| PT2 | 1260 | 69 | 400 ± 120 | 7.9 ± 2.4 | 0.19 ± 0.054 |
| PT3 | 306 | 43 | 150 ± 12 | 2.9 ± 0.25 | 0.3 ± 0.024 |
| PT4 | 288 | 99 | 170 ± 15 | 3.5 ± 0.3 | 0.57 ± 0.049 |
| PT5 | 193 | 39 | 88 ± 12 | 1.8 ± 0.24 | 0.45 ± 0.062 |
| PT6 | 181 | 91 | 120 ± 3.7 | 2.5 ± 0.074 | 0.73 ± 0.022 |
| PT7 | 380 | 8 | 120 ± 14 | 2.5 ± 0.28 | 0.065 ± 0.0077 |
| PT8 | 234 | 74 | 130 ± 4 | 2.6 ± 0.081 | 0.58 ± 0.018 |
| PT9 | 207 | 46 | 99 ± 13 | 2 ± 0.25 | 0.47 ± 0.057 |
| PT10 | 354 | 51 | 140 ± 16 | 2.7 ± 0.32 | 0.38 ± 0.05 |
| PT11 | 438 | 200 | 270 ± 5.1 | 5.5 ± 0.1 | 0.73 ± 0.014 |
| PT12 | 396 | 17 | 120 ± 23 | 2.3 ± 0.45 | 0.15 ± 0.03 |
| PT13 | 233 | 140 | 170 ± 4.1 | 3.4 ± 0.082 | 0.81 ± 0.02 |
| PT14 | 428 | 67 | 180 ± 24 | 3.6 ± 0.47 | 0.38 ± 0.055 |
| PT15 | 180 | 51 | 96 ± 8.5 | 1.9 ± 0.17 | 0.54 ± 0.048 |
| PT16 | 198 | 47 | 96 ± 5.6 | 1.9 ± 0.11 | 0.49 ± 0.027 |

**Table S4.** . Mutated genes restricted to VTT and occurring in more than one patient.

| symbol | patients |
| --- | --- |
| MUC6 | PT11,PT15,PT6 |
| TTN | PT12,PT2,PT7 |
| CFAP54 | PT5,PT8 |
| COL4A3 | PT9,PT7 |
| EIF3FP1 | PT3,PT11 |
| FHAD1 | PT7,PT13 |
| FRG1DP | PT8,PT5 |
| GOLGA6L2 | PT3,PT15 |
| GRIK4 | PT9,PT11 |
| IGFN1 | PT16,PT11 |
| KRTAP5-8 | PT15,PT3 |
| MUC4 | PT7,PT3 |
| NBPF12 | PT2,PT3 |
| PCDH18 | PT3,PT12 |
| PCDHGA3 | PT7,PT14 |
| PRB4 | PT9,PT8 |
| SETD2 | PT6,PT9 |
| SOX9 | PT13,PT12 |
| TSPEAR | PT4,PT2 |
| TYW1 | PT2,PT3 |

**Table S5.**
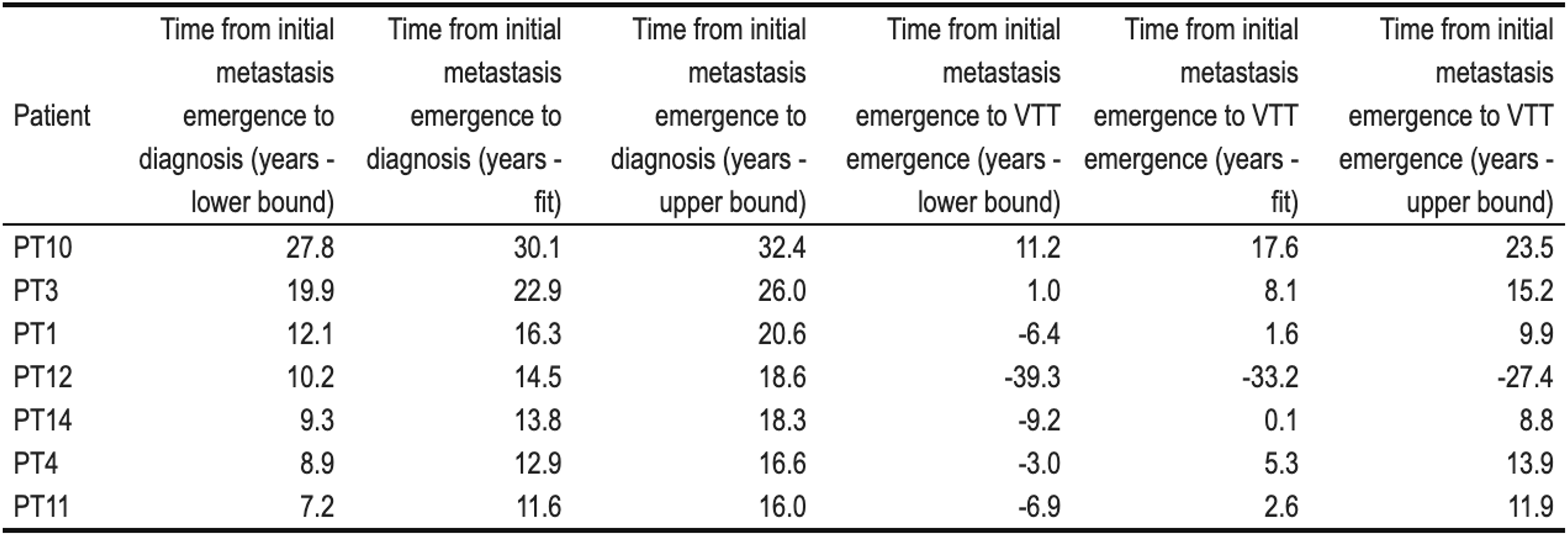
Timing estimates of metastases, VTT, and diagnosis.

